# Dynamic sound field audiometry: static and dynamic spatial hearing tests in the full horizontal plane

**DOI:** 10.1101/849836

**Authors:** T. Fischer, M. Kompis, G. Mantokoudis, M. Caversaccio, W. Wimmer

**Affiliations:** Department of ENT, Head and Neck Surgery, Inselspital, Bern University Hospital, University of Bern, Bern 3008, Switzerland; Hearing Research Laboratory, ARTORG Center for Biomedical Engineering Research, University of Bern, Bern 3008, Switzerland

**Keywords:** sound source tracking, minimum audible angle, auditory motion perception, spatial hearing

## Abstract

Although spatial hearing is of great importance in everyday life, today’s routine audiological test batteries and static test setups assess sound localization, discrimination and tracking abilities rudimentarily and thus provide only a limited interpretation of treatment outcomes regarding spatial hearing performance. To address this limitation, we designed a dynamic sound field test setup and evaluated the sound localization, discrimination and tracking performance of 12 normal-hearing subjects. During testing, participants provided feedback either through a touchpad or through eye tracking. In addition, the influence of head movement on sound-tracking performance was investigated. Our results show that tracking and discrimination performance was significantly better in the frontal azimuth than in the dorsal azimuth. Particularly good performance was observed in the backward direction across localization, discrimination and tracking tests. As expected, free head movement improved sound-tracking abilities. Furthermore, feedback via gaze detection led to larger tracking errors than feedback via the touchpad. We found statistically significant correlations between the static and dynamic tests, which favor the snapshot theory for auditory motion perception.

## 1 Introduction

Reliable localization and tracking of sound sources facilitates speech comprehension in difficult listening situations and is essential for personal safety, for example, in urban traffic scenarios (Cherry 1953). In audiology, sound localization tests are employed to derive binaural information integration ability from spatial hearing performance (Blauert 1997; Middlebrooks 2015).

Thus far, sound field localization experiments with loudspeaker setups are considered the most suitable technique to provide realistic but controlled test conditions, especially with subjects wearing hearing aids and audio processors (Van den Bogaert et al. 2006; Wimmer et al. 2017; Zirn et al. 2019; Denk et al. 2019). Traditionally, tests for spatial hearing in clinics are performed with a static speaker setup covering the frontal azimuth (Grantham et al. 2007; Kerber & Seeber 2012; Dorman et al. 2016). The absolute sound localization tests performed with these setups are in stark contrast to reality, where we are constantly surrounded by dynamic auditory environments with moving sound sources and events. More sophisticated tests involving the perception of auditory movement require a real or simulated displacement of at least one sound source (Grantham 1997; Brimijoin & Akeroyd 2014; Moua et al. 2019). While simulated moving loudspeakers avoid artificial movement noise, which can provide unwanted cues, it remains unclear to what extent they can reproduce physically moving sound sources under consideration of the requirements for clinical testing. To overcome the need for spacious loudspeaker setups for sound localization testing, researchers try to approximate head-related transfer functions in a reasonable time (Mueller et al. 2012; Prepelită et al. 2016; Zhang et al. 2017; Geronazzo et al. 2019). For the creation of complex and dynamic auditory scenes, progress has been made in the area of phantom sound generation techniques (Zhang et al. 2017). However, further investigations are necessary to verify the stability and reproducibility requirements for clinical application, particularly their sensitivity to the listener’s head pose and room characteristics (Zhang et al. 2017; Nelson et al. 2019; ISO 2009).

Due to the rapid progress in the field of computational auditory scene analysis, it is necessary to be able to reliably test and adjust existing and future features of audio processors for clinical purposes (Virtanen et al. 2018). These features include fixed and adaptive beamformers as well as environmentally aware systems such as feature-based classifiers, noise-suppression algorithms, and source-tracking and separation techniques. Further advances can be expected from data-driven artificial intelligence approaches (Naithani et al. 2017; Martinez et al. 2019).

To address these limitations, we developed a robotic measurement setup intended for dynamic sound field audiometry in the azimuthal plane. The system features multiple physically movable loudspeakers with real-time controllable angular position and velocity. Our main objective was to evaluate the possibilities and limitations of the setup by performing a series of spatial hearing experiments in the horizontal plane. The sound localization, discrimination and tracking performance of normal-hearing subjects were measured, and their interrelationships were investigated. The secondary objectives were to assess the influence of head movement on subjects’ sound source tracking performance and to collect reference data on normal-hearing subjects.

## 2 Methods

### 2.1 Study design and participants

This prospective, single-center study was designed in accordance with the Declaration of Helsinki and was approved by our local institutional review board (KEK-BE, No. 2018-00901). Twelve normal-hearing adults between the ages of 24 and 54 years participated in the study^1^. The participants had air conduction hearing thresholds equal to or better than a 15-dB hearing level at 0.5, 1, 2 and 4 kHz. None of the participants were familiar with sound localization studies. The participants gave written informed consent before undergoing the study procedure.

### 2.2 Measurement setup

All test procedures were performed in a sound-attenuated acoustic chamber (6 × 4 × 2m^3^) with an approximate reverberation time of 200 ms for frequencies between 0.25 and 10 kHz. A horizontal circular rail with a radius of 1.1 m was installed at the center of the chamber. Depending on the test procedure, the rail could be equipped with up to 12 loudspeakers (Control 1 Pro, JBL, Northridge, USA). The entire test setup is shown in Figure 1. The loudspeakers were either positioned at a fixed azimuth by using passive support slides or set to arbitrary positions by using wireless controllable audio robots (WCARs). When mounted on the rail, the center of each loudspeaker membrane was at a height of 1.2 m. While the passive slides remained in their initial azimuthal position, a WCAR could be used to present acoustic stimuli at an arbitrary azimuthal position during movement or at rest. The study participants were seated at the center of the circular setup. The setup was controlled using a personal computer and custom-developed software written in C++. The graphical user interface (GUI) was implemented using the QT framework (version 4.2.0, The Qt Company, Espoo, Finland). User feedback was provided either by a touchscreen showing test-specific GUIs or by gaze detection. Audio processing was performed with the FMOD engine (Firelight Technologies Pty Ltd, Melbourne, Australia). Using the software, various test parameters could be defined and scripted to sequences and trajectories, including the azimuthal position and angular velocity of the WCAR as well as the level, duration and type of acoustic stimuli. The wireless data transfer between the operating station and each WCAR was realized via a proprietary wireless local area network protocol.

**Figure 1:**
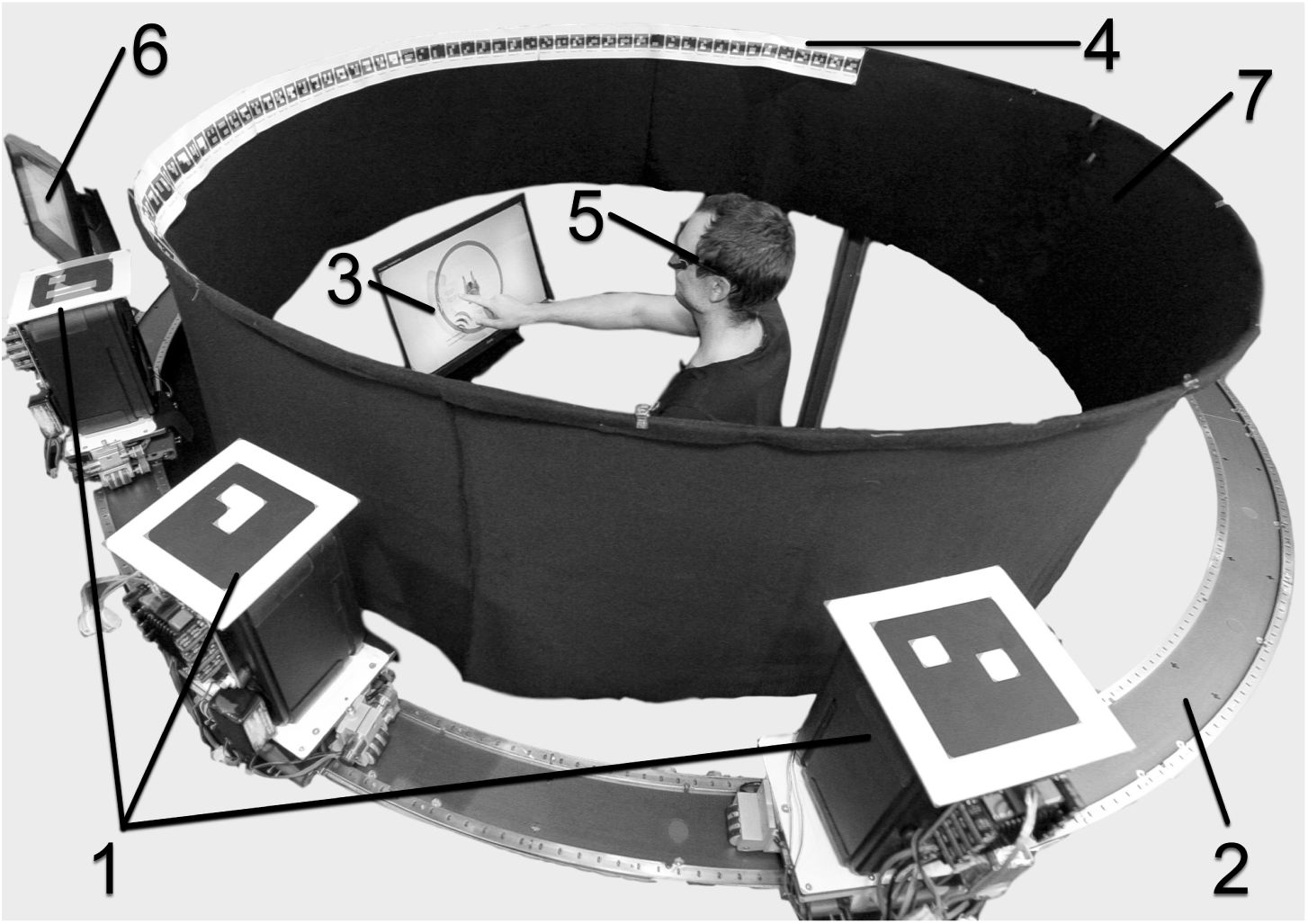
The robotic measurement setup during minimum audible angle (MAA) assessment at 270 degrees azimuth. Three wireless controllable audio robots (WCARs) with optical tracking markers (1), low noise rail (2), touch screen with graphical user interface (GUI) (3), eye-tracking glasses (4), azimuth reference markers (5), control terminal (6) and sound-transparent curtain (7).

Each WCAR contained a single-board computer (Raspberry Pi 3 Model B, RASPBERRY PI FOUNDATION, Caldecote, UK) to enable wireless communication and on-board signal processing. The board was extended by a 192-kHz/24-bit audio output shield (HiFiBerry DAC+, Modul 9 GmbH, Ermatingen, Switzerland). To drive the loudspeakers, 200 W class D amplifiers (AA-AB31282, Sure Electronics, Nanjing, China) were used. The WCARs were positioned using a DC motor and a dedicated motor controller (DCX 19S and EPOS2 24/2 DCX, maxon motor AG, Sachseln, Switzerland). The power transmission for WCAR movement consisted of a rubber-toothed belt connected to a rubber roller. The mechanical components were designed to minimize movement noise. At the maximum velocity that occurred in our test procedures (7.4°/s corresponding to 0.14 m/s), a moving WCAR produced movement noise with a 34.9-dB sound pressure level (SPL) (root mean squared average between 100 Hz and 10 kHz)^2^.

Power management of the on-board computer was controlled by a supply circuit and battery (PiJuice 3.7 V, 1820 mAh, Pi-Supply, Kent, United Kingdom). In addition, the audio power amplifier and the motor controller were each supplied by separate lithium-polymer batteries (14.5 V and 18.5 V, 1300 mAh). Position tracking of the WCARs was performed using 2 redundant systems. First, the position encoder of the motor was continuously read out by the single-board computer at a 5 Hz sampling rate. Second, an optical tracking system consisting of 3 cameras (USBFHD06H-BFV, ELP, China) and fiducial markers specifying each loudspeaker individually (ArUco Framework) was applied (Romero-Ramirez et al. 2018). The combination of the motor encoder and the optical tracking enabled a loudspeaker positioning accuracy of 0.5°.

Each loudspeaker was equalized and calibrated with a free-field microphone (type 4133 and preamplifier type 2639, Brüel & Kjaer, Nærum, Denmark) positioned at the reference point in the center of the circular setup and an audio analyzer (UPV, Rohde & Schwarz, Munich, Germany). The calibration data of each loudspeaker were stored individually to achieve a flat frequency spectrum between 0.15 and 10 kHz. The loudspeakers were covered by a sound-transparent black curtain to avoid visual cues during the localization tasks.

### 2.3 Head orientation and eye tracking

During the entire session, the participants wore eye-tracking glasses equipped with a 120-Hz world camera and 200-Hz binocular eye cameras (Pupil Labs, Berlin, Germany). The data of the eye-tracking glasses were used to monitor the head and gaze positions of the participants during the tests to minimize orientation-based systematic errors (Razavi et al. 2007). Gaze tracking was used as an additional response method for the sound source tracking tests. The light conditions were kept constant during all measurements. To minimize bias in pupil and gaze detection, no light sources were inside the visual field of the subjects (Winn et al. 2018). In the frontal azimuth (i.e., 0° ± 90° azimuth), fiducial markers with labels indicating their corresponding azimuth were attached to the sound-transparent curtain. In total, 61 markers were placed with an angular spacing of 3° (see Figure 1). The markers were used to calculate the head orientation and gaze direction using a Python script (Pupil Player v1.10, Pupil Labs).

### 2.4 Test procedures

Each subject participated in a session consisting of the following 4 blocks: (1) static sound source localization for testing absolute sound localization, (2) MAA assessment for testing sound discrimination, (3) sound source tracking with GUI, and (4) sound source tracking with eye and head tracking for testing dynamic tracking abilities and the effects of head movements on these abilities. Each block was introduced through a training. If desired by the participants, short breaks were taken in between blocks. To minimize systematic training and fatigue effects, the block sequence was counterbalanced. In addition, counterbalancing of specific test conditions within a block was performed if applicable (as specified for each test procedure). During testing, the head orientation of the participants was monitored, as described in subsection 2.3, to ensure a central head position and a correct alignment of the interaural axis.

#### 2.4.1 Static sound source localization

Static sound source localization was assessed using 12 loudspeakers at fixed positions arranged in the full azimuthal plane, resulting in a 30° angular spacing. Pink noise 200 ms in duration with levels ranging between 60 and 70 dB SPL was used as the acoustic stimulus. Three stimuli per loudspeaker were presented in a randomized order, resulting in a total of 36 stimuli per test (Wimmer et al. 2017). The participants were instructed that stimuli could be presented from any azimuth. They indicated the location of the stimulus source on a touch interface with a 1° resolution dial. Two seconds after the subjects logged in their answer, the next stimulus was triggered. Before testing, the participants trained the input method until they confirmed that they fully understood the test procedure. No feedback about the performance was provided during or after testing. A Monte Carlo simulation yielded a chance level of RMSE_LOC_ = 103.6°.

#### 2.4.2 Minimum audible angle

The MAA is defined as the smallest reliably discriminated change in the angular displacement of a sound source (Mills 1958). We measured the MAA at 0°, 45°, 90°, 135°, 180°, 225°, 270° and 315° azimuths for each participant. First, a 200-ms pink noise pulse with a level of 65 dB SPL was emitted from the reference position. After a 1-s intra-stimulus interval, a second identical impulse was emitted, but it was displaced in either clockwise or counterclockwise directions. Each new step size resulted in an update of the WCAR positions. For small displacements, only one WCAR was used for the reference and displaced stimulus presentation because two WCARs could not go closer than 18.5° due to their size. The movement was timed to ensure a 1-second intra-stimulus duration independent of the size of the displacement. Noise was masked by the simultaneous movement of the 2 other WCARs with an angular offset of 120° each. With this setting, it was not possible to localize the individual movement of speakers. The participants were asked to indicate the direction of the stimulus shift directly via the touch pad^3^. This method was chosen to avoid confusion regarding direction when testing was performed in the area behind the subjects. No feedback was provided during or after the test procedures. The first measurement position was always at the front at 0°. To minimize bias, the remaining 7 measurement positions were tested in either clockwise or counterclockwise order. Due to this counterbalancing, the second measurement position was either at 45° or at 315°. Monte Carlo simulation of the test showed a mean chance performance of 82.6°.

#### 2.4.3 Sound source tracking with touch pad

To test sound source tracking abilities, a single WCAR was mounted on the horizontal rail behind the sound-transparent curtain. The WCAR moved along a predefined trajectory while a pink noise stimulus was continuously played at a level of 65 dB SPL. The participants were instructed to indicate the position of the WCAR on a touch pad displaying a dial with 1° resolution^4^. The input sampling rate of the indicated azimuth was downsampled to 5 Hz to match the sampling rate of the WCAR position encoded in the single-board computer. The angular velocity and the direction of motion were obtained from the first derivative of the user input signal. Small and fast changes in user input are considered artifacts; therefore, we low-pass filtered the velocity signal with a cut-off frequency of 0.5 Hz.

To familiarize the participants with the input method, a training trial covering 450° in azimuth with a single change in direction (CID) at 45° was performed. During the training trial, the real-time position of the loudspeaker was additionally indicated on the GUI as a reference. Two different trajectories covering the whole azimuthal plane were used for testing^5^. Videos of the performed trajectories are provided as multimedia material ^6 7 8^. The first trajectory (steady trajectory) covered a total of 900° starting at 315° with a single CID at 45°. The second trajectory (alternating trajectory) covered a total of 2070° starting at 0° with 32 CIDs each at a multiple of 45°, where each CID position was approached 2 times clockwise and 2 times counterclockwise. The maximum angular velocity was 7.4°/s for both trajectories. This velocity was chosen based on previous studies suggesting an optimal angular velocity for movement detection between 1°/s and 20°/s when using broadband stimuli (Kourosh & Perrott 1990). The WCARs needed 200 ms to accelerate to maximum velocity or to decelerate from maximum speed to standstill. Special care was taken to avoid visual cues that could bias the responses of the participants (e.g., reflections or shadows).

#### 2.4.4 Sound source tracking with/without head movement

To evaluate the influence of head movement on sound source tracking accuracy, we additionally performed a set of tracking experiments in which the participants were instructed to follow the sound source with their eyes. Eye-tracking glasses were used for automated gaze detection. All participants confirmed that they could clearly see the contours of the azimuth reference markers (see Figure 1). The accuracy of the calibrated eye-tracking glasses averaged over all participants was 2.6° (± 1.0°). For a first trial run and for eye-tracking calibration validation, the curtain was lifted to make the moving WCAR visible to the participants. In addition to the calibration procedure, the smooth-pursuit movements and oculomotor function were evaluated by visual inspection of the gaze points during the trial run. For testing, two different trajectories were set up: a steady trajectory (total coverage of 720° with 5 CIDs) and an alternating trajectory with several short movements (total coverage of 885° with 19 CIDs). Both trajectories were limited to the visual field, i.e., 0° ± 60° (Trattler et al. 2012). A a tabular description of the used trajectories is provided in the supplementary materials^9^. Videos of the performed trajectories are provided as multimedia material ^10 11 12^. The tracking experiments were repeated in two conditions. In the first condition, the participants were instructed to keep their head straight while following the sound source with their eyes. In the second test condition, the participants were allowed to freely move their head while following the sound source. We discarded all gaze data with a detection confidence of less than 60%, as recommended by the manufacturer. The remaining data points were interpolated to a sampling rate of 5 Hz to enable a direct comparison with the position data of the WCAR and the GUI input. Unacceptably large data points gaps with separation greater than 500 ms were labelled missing. Subjects 1 and 11 were completely excluded from the analysis due to an overall low detection confidence of their gaze data. Since the data quality varied with respect to the azimuths, all gaze-based analyses used data points for which at least 80% of participants provided valid data.

### 2.5 Outcome measures

#### 2.5.1 Static sound source localization

To assess the absolute localization accuracy, we computed the root mean square error (RMSE) between the azimuthal positions of the stimulating loudspeakers *ϑ*_*n*_ and the indicated positions 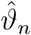 (in degrees), averaged over the total number stimuli (*N* = 36) (Hartmann 1983):

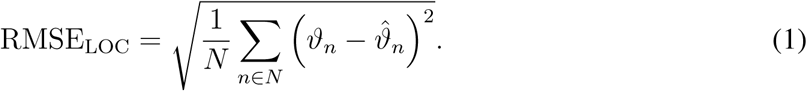

As recommended, front-back confusions (FBCs) were excluded from the errors and analyzed separately (Letowski & Letowski 2011).

#### 2.5.2 Minimum audible angle

Each MAA step size was determined following an updated maximum-likelihood estimation assuming a logistic-shaped psychometric function with a start measurement distance of 15° (Shen et al. 2015). The test was implemented as a 2-alternative forced choice procedure applying a 2-down, 1-up rule to minimize the impact of guessing. The parameter range of the angular displacement was set from 0.5° to 90°. For each measurement position, 24 measurements with an adaptive step size were performed. The MAA value was defined as the 80%-correct threshold of the estimated psychometric function.

#### 2.5.3 Sound source tracking with touch pad

During the tracking tests the participants continuously indicated the angular position of the moving sound source. Similar to the static source localization test, we assessed the tracking accuracy using the RMSE. The subject input 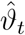 and actual WCAR position *ϑ*_*t*_ (in degrees) sampled at time *t* and averaged over the test duration *T* was calculated:

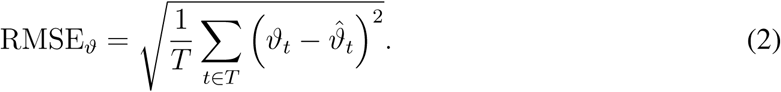

To quantify the tracking velocity accuracy between user input 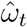 and actual angular velocity *ω*_*t*_ we used:

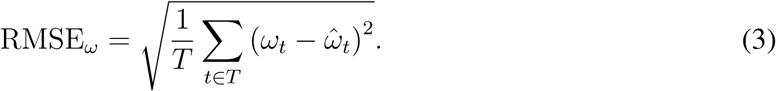

To exclude effects caused by individual reaction times, the first 4 seconds after stimulus onset were discarded from the analysis. The percentage of congruence between the indicated and actual directions of motion (i.e., counterclockwise vs. clockwise) was computed for the total procedure

To measure the sensitivity to a change in direction (CID), the elapsed time and azimuthal position were recorded until the participant adapted to the change. The following criteria had to be fulfilled for a CID to be classified as valid: (i) the movement direction before the change was correctly indicated by the subject, (ii) the subject could follow the correct direction after the CID for at least 2 seconds, and (iii) the subject perceived the change within 20 s. For each CID, we recorded time delay Δ*t*_cid_ between actual change *t*_cid_ and the time indicated by the participant, 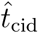 (in ms), as well as the angular displacement Δ*ϑ*_cid_ (in degrees) that occurred at 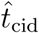 (see Figure 2).

**Figure 2:**
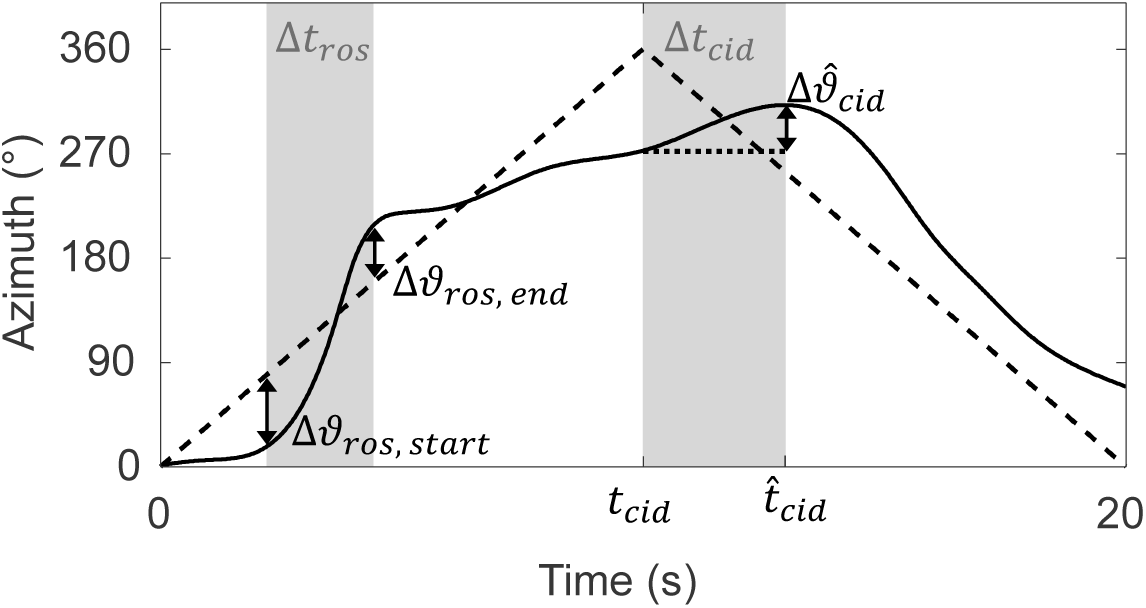
Example trajectories of a moving stimulus (dashed line) and the corresponding user input (solid line). A re-localization of source (ROS) event, in which the indicated velocity exceeds 3 times the angular velocity of the moving sound source, is shown. The ROS has a duration of Δ*t*_ros_. A change in direction (CID) at 360° azimuth occurs after *t*_cid_ = 10 seconds. The time it took the subject to indicate the change (Δ*t*_cid_) and the azimuthal offset 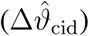 were recorded.

Whenever the indicated angular velocity was 3 times higher than the velocity of the moving sound source, we assumed that the participant lost track of the stimulus and had to re-localize it. We refer to such an event as a re-localization of source (ROS). We recorded the number of ROS occurrences *N*_ros_ as well as the time Δ*t*_ros_ it took the participant to re-localize the sound source (in ms). The error before Δ*ϑ*_ros,start_ and after a sound search Δ*ϑ*_ros,end_ was calculated with ΔRMSE_*ϑ*,ros_ (in °).

#### 2.5.4 Head movement and tracking accuracy

To evaluate the contributions of head orientation and gaze direction to the indication of the stimulus position, we compared the position and velocity-tracking RMSEs between the two conditions (head movement restricted vs. head movement unconstrained). In addition, the congruence between the head orientation and the angular sound source position was quantified with the Pearson correlation coefficient.

#### 2.5.5 Comparison touch pad vs. gaze detection input

To evaluate the differences between the GUI-based and gaze detection-based input methods, we compared the data obtained during the test condition of restricted head movement with participants’ corresponding GUI-based input. The greatest advantage of the eye-tracking method is that it represents an immediate response behavior; however, it is limited to participants’ visual field when their head movement is restricted. Even when head and torso movements are unconstrained, it is difficult to cover the whole azimuth (especially the area behind participants) in a convenient way. For this reason, we used only data from the two input methods within the range of 0° ± 22.5°. To evaluate the differences in the input methods, we compared the reaction times of the participants after a frontal CID. In addition, we compared the RMSE_*ϑ*_ of both methods within the tested azimuth. Five of 12 subjects were excluded from the analysis due to insufficient data quality.

### 2.6 Statistical analysis

Descriptive statistics were used to quantify the performance of the subjects on a group level. To compare localization and tracking errors under different conditions a two-tailed Wilcoxon signed rank test was used. The Pearson correlation coefficient was used to investigate relationships between the different test procedures and outcome measures at the subject level. A significance level of *α* = 0.05 was chosen. Statistical calculations were performed with Matlab (version R2018a, The MathWorks Inc, USA) and the CircStat toolbox (Berens 2009).

## 3 Results

### 3.1 Static sound source localization

Figure 3 illustrates the absolute localization accuracy for each azimuth under exclusion of FBCs. Participants’ performance was best at the front and the back, with average RMSE_LOC_ values of 3.9° (standard deviation, ± 3.4°) and 6.2° (± 5.6°), respectively (p>0.05). The participants had a better localization accuracy on the right side (7.8° ± 7.4°) than on the left side (15.9° ± 9.1°) (p<0.05). Excluding the left and right directions, the localization performance in frontal azimuths (11.8° ± 3.9°) was comparable to that in dorsal azimuths (13.8° ± 4.8°) (p>0.05). The overall RMSE_LOC_ was 12.9° ± 5.9°. The mean response time for each stimulus was similar across directions, with an average of 6.60 s (±0.25 s). We did not observe any correlation between the response time and localization performance. The individual performance of the participants is shown in the supplementary materials^13^. Five subjects had FBCs resulting in an FBC rate of 3.6%. Of the total of 13 FBCs, 6 were back-front confusions.

**Figure 3:**
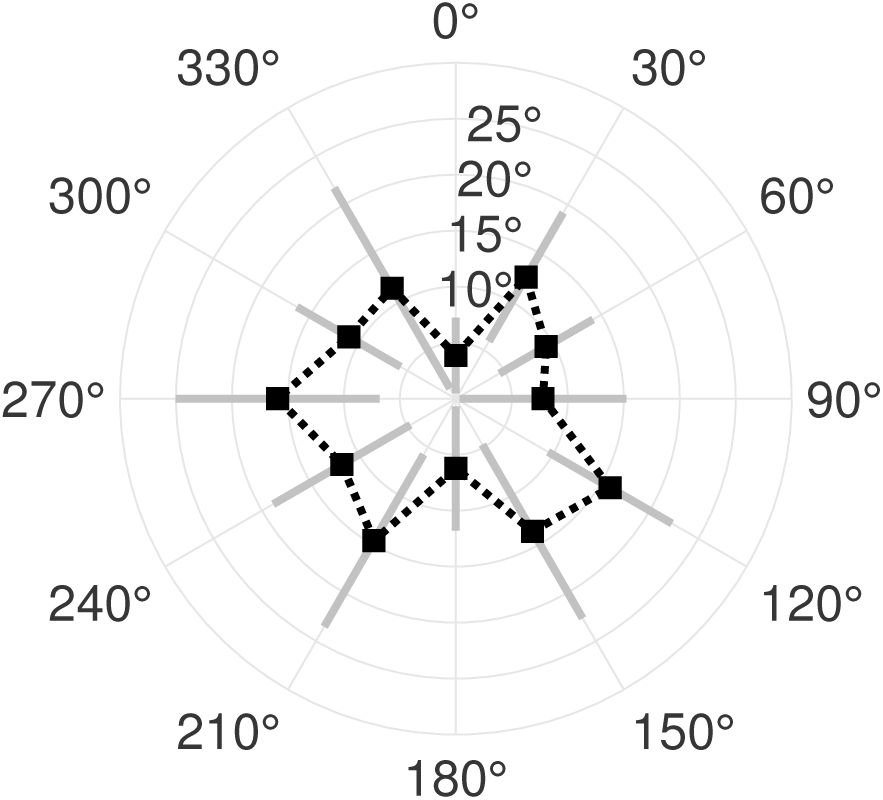
Root mean square localization errors (RMSE_LOC_) averaged across all participants for each stimulus direction in the static sound source localization test. Gray bars indicate one standard deviation.

### 3.2 Minimum audible angle

MAA performance was best at the front with 1.2° (±0.4°) and worst at the right and left sides with 6.5° (±4.2°) and 4.8° (±2.7°), respectively. A polar plot of the MAA results averaged across all participants is shown in Figure 4. Subject- and azimuth-specific MAA results are listed in the supplementary materials^14^.

**Figure 4:**
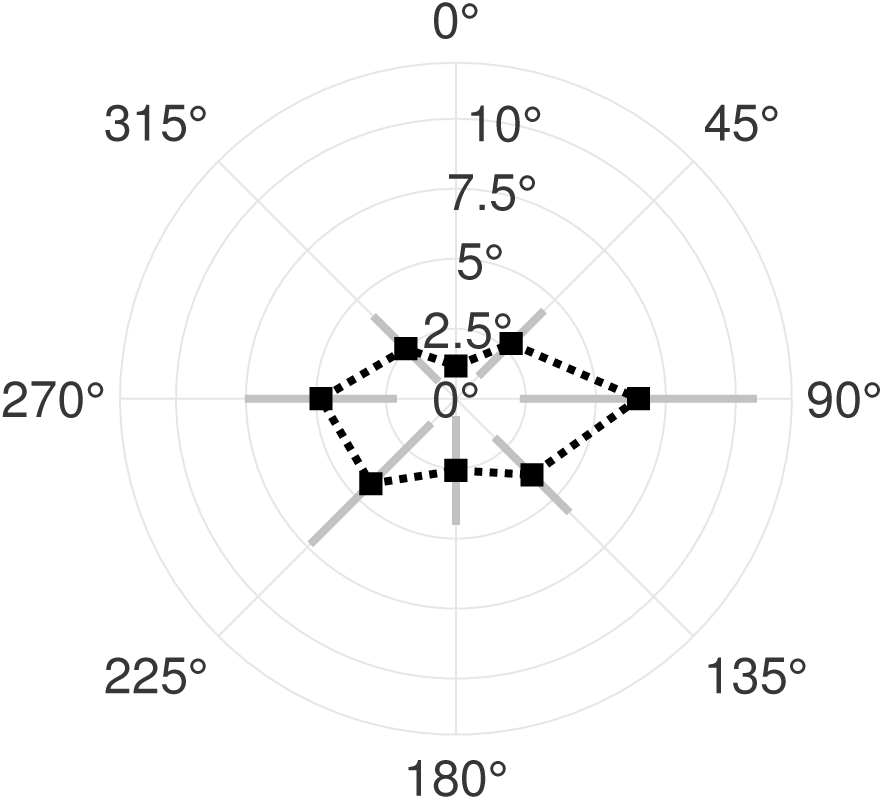
Minimum audible angle (MAA) performance averaged across all participants with respect to the tested azimuth. Gray bars indicate one standard deviation.

Excluding the left and right directions, the MAA in frontal azimuths (2.2° ± 1.0°) was significantly lower than that in dorsal azimuths (3.6° ± 1.8°) (p<0.05). Without the positions at 0° and 180°, the MAA on the left side (3.9° ± 1.8°) was comparable to that on the right side (4.4° ± 1.8°) (p>0.05). The overall MAA averaged over all subjects was 3.6° ± 1.4°.

### 3.3 Sound source tracking with touch pad

A summary of the sound source tracking performance for the steady (Video MM2.avi) and alternating trajectory (Video MM3.avi) is provided in Table 1. Figure 5 illustrates the position of the WCAR, the GUI-based input and the tracking errors during the steady trajectory experiment. Most of the time, the participants indicated the movement directions correctly and had no problems following the sound source. The mean error shows that the participants tended to indicate the position of the sound source ahead of the stimulus, independent of the movement direction.

Most prominently, a response delay occurred following the single CID at 45° azimuth causing the largest tracking errors (RMSE_*ϑ*_ of 31.5° averaged across all tested subjects). The smallest tracking errors were observed towards the front (0°) and the back (180°). Figure 6 shows the position of the WCAR and the corresponding GUI-based input with the source tracking errors for the alternating trajectory tests. The overall position tracking error (RMSE_*ϑ*_) was 25 % lower in the tests with the steady trajectory than in those with the alternating trajectory. Similar to the steady trajectory case, delays in position tracking after each CID were observed, with an average delay (Δ*t*_cid_) of 1.3 seconds. In Figure 7, a polar representation of the tracking errors is plotted for both movement directions (clockwise versus counterclockwise) during the alternating trajectory experiment. The subjects had the same performance in correctly following sound sources moving in a counterclockwise direction (RMSE_*ϑ*_ of 18.9°± 4.9°) and in a clockwise direction (RMSE_*ϑ*_ of 18.4°± 5.5°, p=0.7). The difference was attributed more to larger delays directly after CIDs and less to tracking offsets outside of CIDs (Figure 7). The RMSE_*ϑ*_ peaks were shifted by delay Δ*t*_*cid*_ in the corresponding movement direction. A direction specific plot of the CID reaction times is shown in Figure 9. The smallest tracking errors occurred at frontal azimuths for either movement direction at 0°± 45°. The largest errors were observed after the sound source crossed the interaural axis in dorsal directions. Overall, the participants had a higher position tracking accuracy in the frontal azimuths than in the dorsal azimuths, regardless of the movement direction (RMSE_*ϑ*_ in clockwise directions of 15.6° ± 3.9° versus 20.7° ± 7.2° and in counterclockwise directions of 17.0° ± 4.6° versus 20.2° ± 6.0°; both p<.05). A similar tracking accuracy for stimuli in the left and right hemifields was achieved. However, for counterclockwise movements, the RMSE_*ϑ*_ was larger in the left hemifield (left hemifield 19.6° ± 5.6° vs. right hemifield 17.9° ± 5.4°, p = 0.30), and for clockwise movements, the error was larger in the right hemifield (16.2° ± 5.5° vs. right hemifield 20.1° ± 6.6°, p = 0.09). Thus, crossing the interaural axis and moving towards the back caused the largestRMSE_*ϑ*_ (see Figure 7).

**Table 1:** Summary of outcome measures for sound source tracking using a touch pad. SD = standard deviation, RMSE = root mean square error, CID = change in direction, ROS = re-localization of source.

|  | Steady trajectory |  | Alternating trajectory |  |
| --- | --- | --- | --- | --- |
|  | Mean | SD | Mean | SD |
| Position tracking error, $\text{RMSE}_{\theta}$ ($^{\circ}$) | 14.1 | 2.0 | 18.7 | 5.0 |
| Velocity tracking error, $\text{RMSE}_{\omega}$ ($^{\circ}/\text{s}$) | 0.9 | 0.1 | 1.6 | 0.4 |
| Correctly indicated direction (%) | 97.2 | 1.6 | 82.5 | 4.8 |
| Delay after CID, $\Delta t_{\text{cid}}$ (ms) | 1'688 | 455 | 1'341 | 439 |
| Angular offset after CID, $\Delta \hat{\vartheta}_{\text{cid}}$ ($^{\circ}$) | 11.7 | 4.1 | 11.4 | 4.9 |
| Number of ROS events, $N_{\text{ros}}$ | 1 | 1 | 14 | 12 |
| ROS duration, $\Delta t_{\text{ros}}$ (ms) | 684 | 678 | 498 | 189 |
| Improvement after ROS, $\Delta \text{RMSE}_{\theta, \text{ros}}$ ($^{\circ}$) | 13.0 | 11.1 | 14.1 | 7.1 |

**Figure 5:**
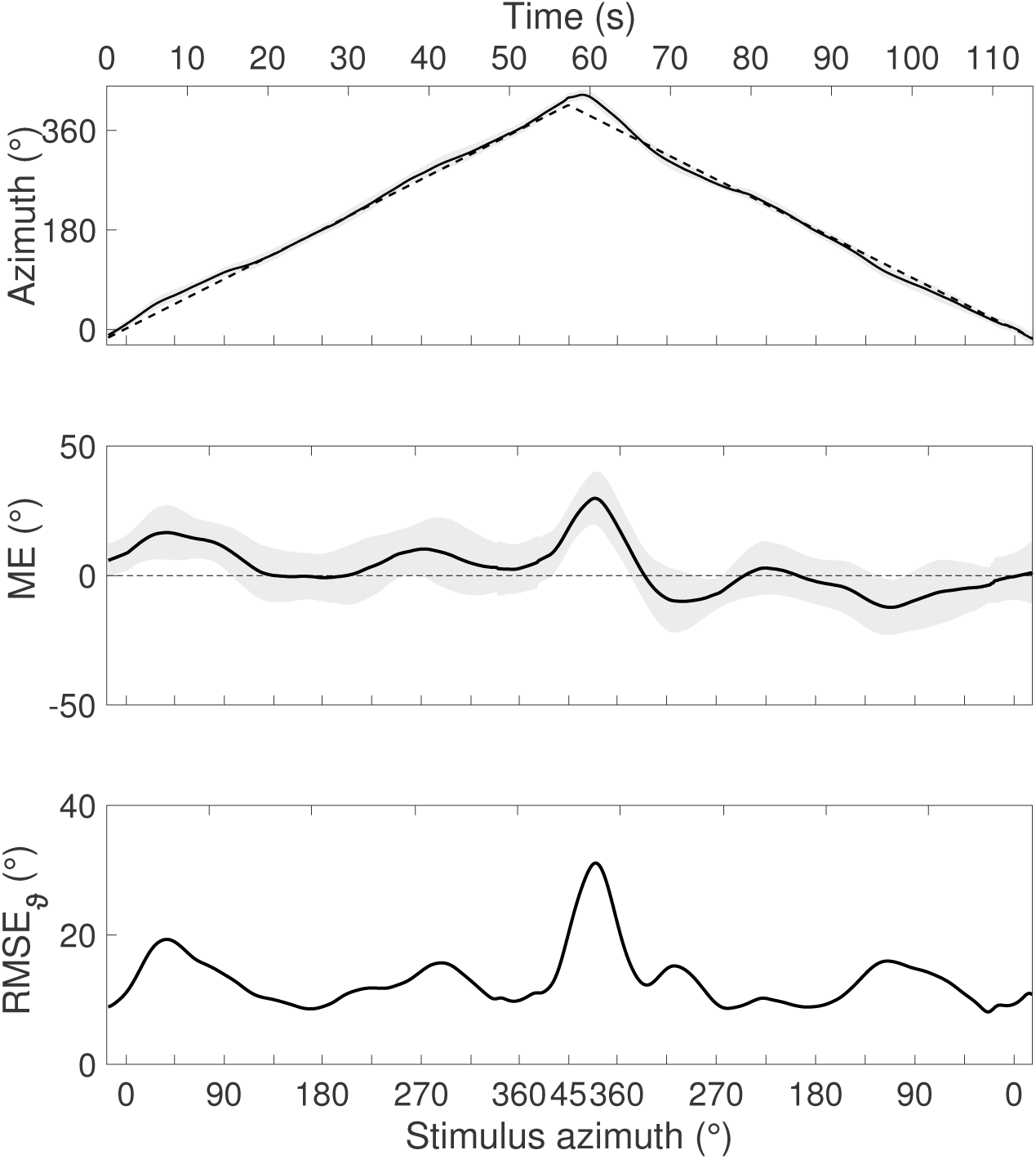
Sound source tracking during the steady movement trajectory with a single change of direction at 45° azimuth. Plotted are the location of the sound source and the averaged subject responses (as indicated with a the touch pad) as well as the mean error (ME) and root mean square error (RMSE_*ϑ*_) averaged across all subjects.

**Figure 6:**
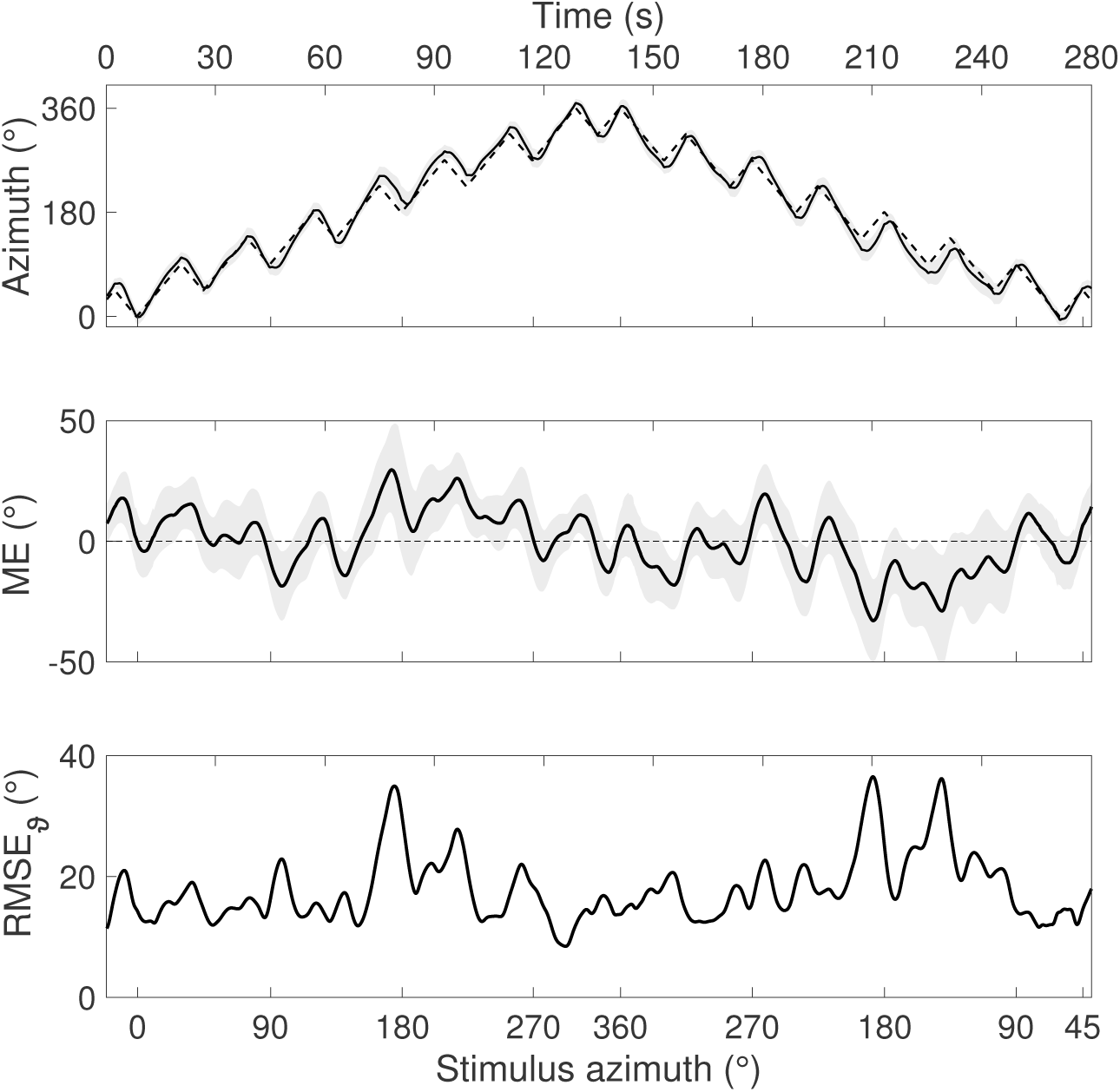
Sound source tracking of the alternating movement trajectory with multiple changes of direction. Plotted are the location of the sound source and the averaged subject responses (indicated with touch pad) as well as the mean errors (MEs) and root mean square error (RMSE_*ϑ*_) averaged across all subjects.

**Figure 7:**
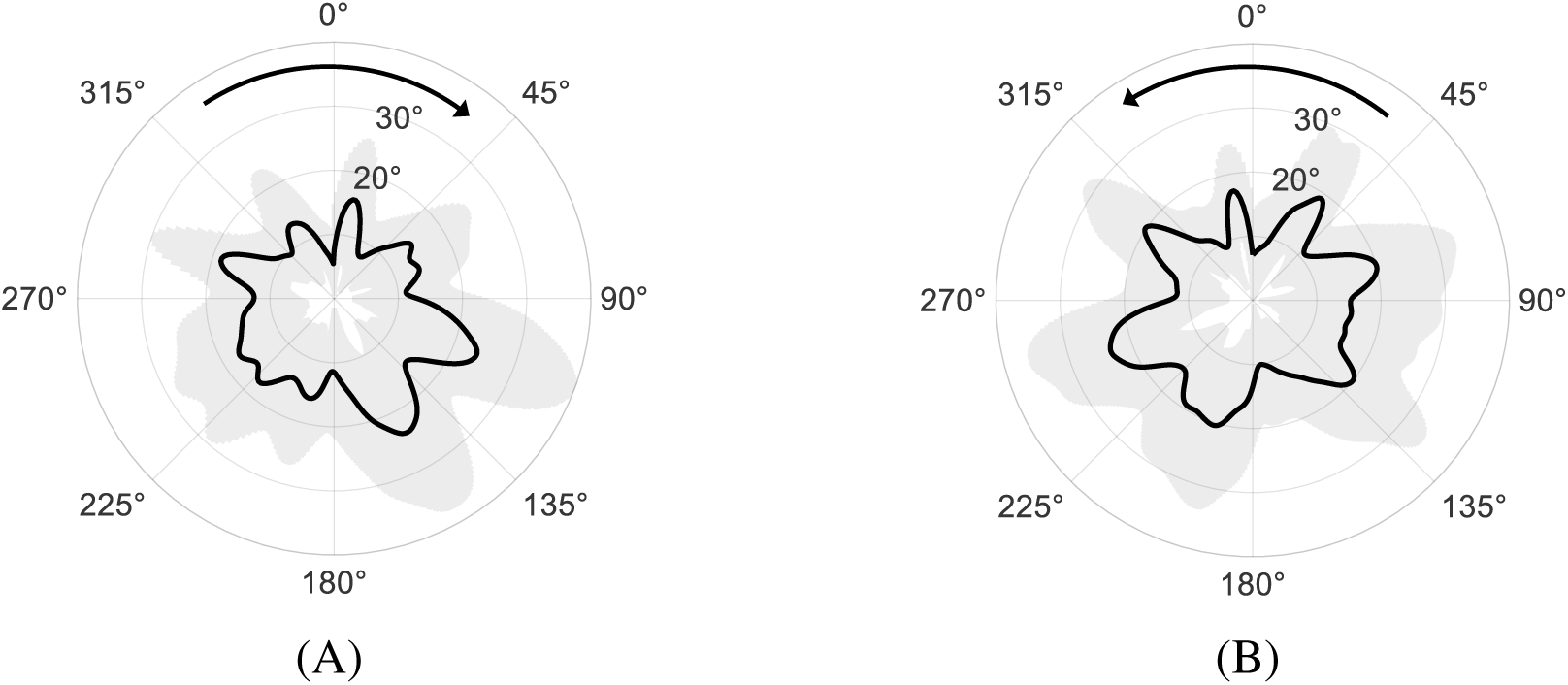
Root mean square error (RMSE_*ϑ*_) averaged over all subjects during the alternating sound source tracking task with the touch pad based feedback method. The errors are shown separately for clockwise (A) and counterclockwise (B) movements of the sound source. Gray areas indicate ±1 standard deviation.

**Figure 8:**
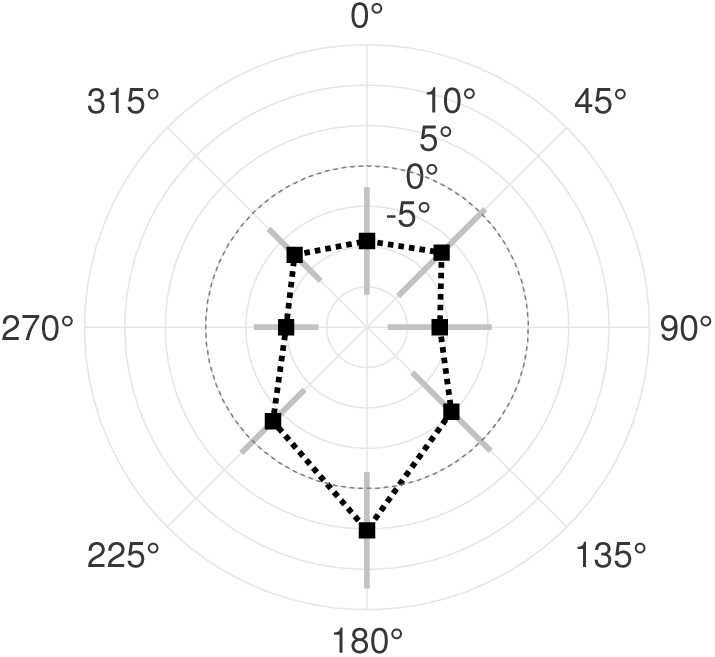
Relative improvement in sound source tracking accuracy 2 s before and after repeated changes in direction (CIDs) at 8 different azimuths. Negative values indicate worse performance after the CID.

**Figure 9:**
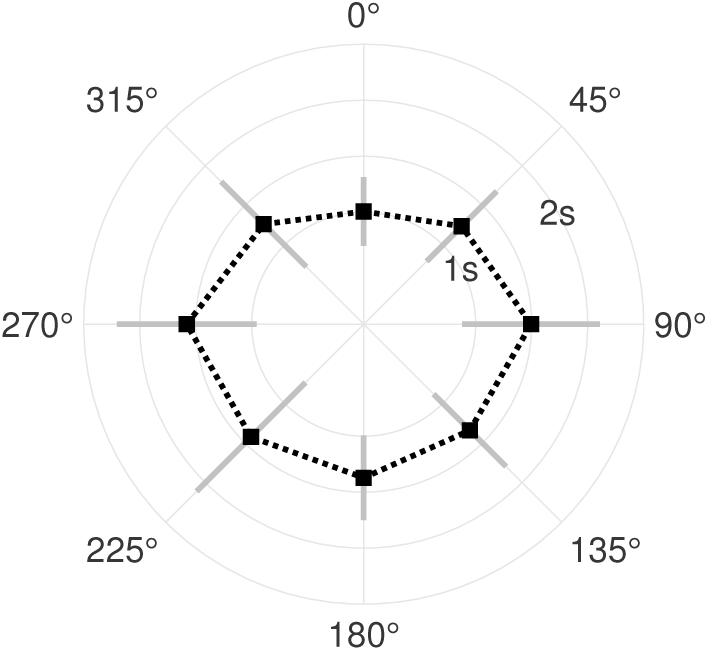
Reaction times taken to indicate a changes in direction (CID) (Δ*t*_cid_) at the different azimuths.

The angular velocity tracking error RMSE_*ω*_ was almost twice as high for the alternating trajectory than for the steady trajectory (Table 1). This difference could be explained by the higher incidences of CIDs during the alternating trajectory tests.

Figure 8 illustrates the relative change in tracking accuracy after the indication of a CID for the corresponding azimuth. With the exception of the position at 180°, the tracking accuracy deteriorated after the change was recognized. At the interaural axis (i.e., at 90° and 270°), CIDs led to the greatest deterioration of position tracking accuracy. At 180° azimuth, an improvement of tracking accuracy after a CID was observed. The fastest responses to CIDs occurred in the front at 315° (1’227 ms ± 541 ms), 0° (964 ms ± 283 ms), and 45° (1’212 ms ± 446 ms) azimuths. In other directions, very similar reaction times with a mean value of 1’353 ms and a small standard deviation of 78 ms were observed (see Figure 9).

The higher tracking uncertainty during the alternating trajectory tests was also reflected in the substantially higher incidence of ROS events (Table 1). Most of the ROS events started after the azimuthal deviation between the stimulus and the response exceeded 10 degrees. No correlation could be observed between the number of ROSs and the azimuth of the sound source. Overall, the participants achieved lower scores on correctly identifying the movement direction in the alternating trajectory case.

### 3.4 Head movement and tracking accuracy

Independent of the tested trajectories (Videos MM5.avi and MM6.avi), the sound source tracking performance was significantly better when the participants were allowed to move their head (Table 2). Although they were not explicitly instructed to follow the sound source with their head (but were instructed to do so with their eyes), a strong correlation between the head orientation and the sound source position was observed for all participants when head movement was allowed.

**Table 2:** Summary of outcome measures for sound source tracking using gaze detection. SD = standard deviation, RMSE = root mean square error, CID = change in direction, ROS = re-localization of source.

|  | Steady trajectory |  | Alternating trajectory |  |
| --- | --- | --- | --- | --- |
|  | Mean | SD | Mean | SD |
| **Gaze position tracking error** |  |  |  |  |
| Without head movement, $\text{RMSE}_\theta$ ($^\circ$) | 35.4 | 18.9 | 34.6 | 15.2 |
| With head movement, $\text{RMSE}_\theta$ ($^\circ$) | 25.4 | 14.6 | 25.4 | 14.3 |
| **Head orientation only** |  |  |  |  |
| Position tracking error, $\text{RMSE}_\theta$ ($^\circ$) | 11.8 | 3.1 | 10.3 | 3.3 |
| Velocity tracking error, $\text{RMSE}_\omega$ ($^\circ/\text{s}$) | 0.8 | 0.2 | 1.0 | 0.1 |
| Correctly indicated direction (%) | 94.0 | 2.9 | 86.9 | 3.4 |
| Pearson correlation coefficient of head orientation with stimulus position | 0.96 | 0.03 | 0.98 | 0.10 |
| Delay after CID, $\Delta t_{cid}$ (ms) | 753 | 282 | 667 | 223 |
| Angular offset after CID, $\Delta \hat{\vartheta}_{cid}$ ($^\circ$) | 2.8 | 1.6 | 2.4 | 1.1 |
| Number of ROS events, $N_{ros}$ | 1 | 1 | 1 | 1 |
| ROS duration, $\Delta t_{ros}$ (ms) | 374 | 93 | 86 | 177 |
| Improvement after ROS, $\Delta \text{RMSE}_{\theta,ros}$ ($^\circ$) | 1.3 | 0.9 | 0.9 | 0.6 |

Due to the numerous CIDs in the alternating trajectory experiment, the percentage of identical movement directions of the stimulus and head was lower in that experiment than in the steady trajectory experiments. The reaction times and velocity RMSEs were approximately 20% higher for the steady trajectory than for the alternating trajectory. In terms of errors, the greatest offsets appeared at the most lateral positions (±60°) due to limited motor skills (no torso movement allowed). For the steady trajectory, the RMSE_*ϑ*_ of the head position was 11.8 ± 3.1, approximately half the size of the corresponding gaze positions, 25.4 ± 14.6. The same relationship applied for the alternating trajectory with an RMSE_*ϑ*_ of 10.3° ± 3.3° for the head position and 25.4° ± 14.3° for the gaze position.

### 3.5 Comparison touch pad vs. gaze detection input

The tracking RMSE_*ϑ*_ was significantly smaller when the touch pad-based input method (12.5° ± 8.6°) was used than when the gaze-based input method (28.1° ± 16.6°) was used. However, the reaction times Δ*t*_*cid*_ at the front were 953 ms ± 212 ms for the GUI and 363 ms±185 ms for the gaze-based input method. The difference between the two input methods was therefore 590 ms (p<0.05). For comparison, the general response times to auditory stimuli while pushing a button are ∼270 ms for women and ∼300 ms for men (Shelton et al. 2010).

### 3.6 Correlation analysis

Figure 10 displays the scatter plots of selected outcome measures for a comparison at the subject level.

**Figure 10:**
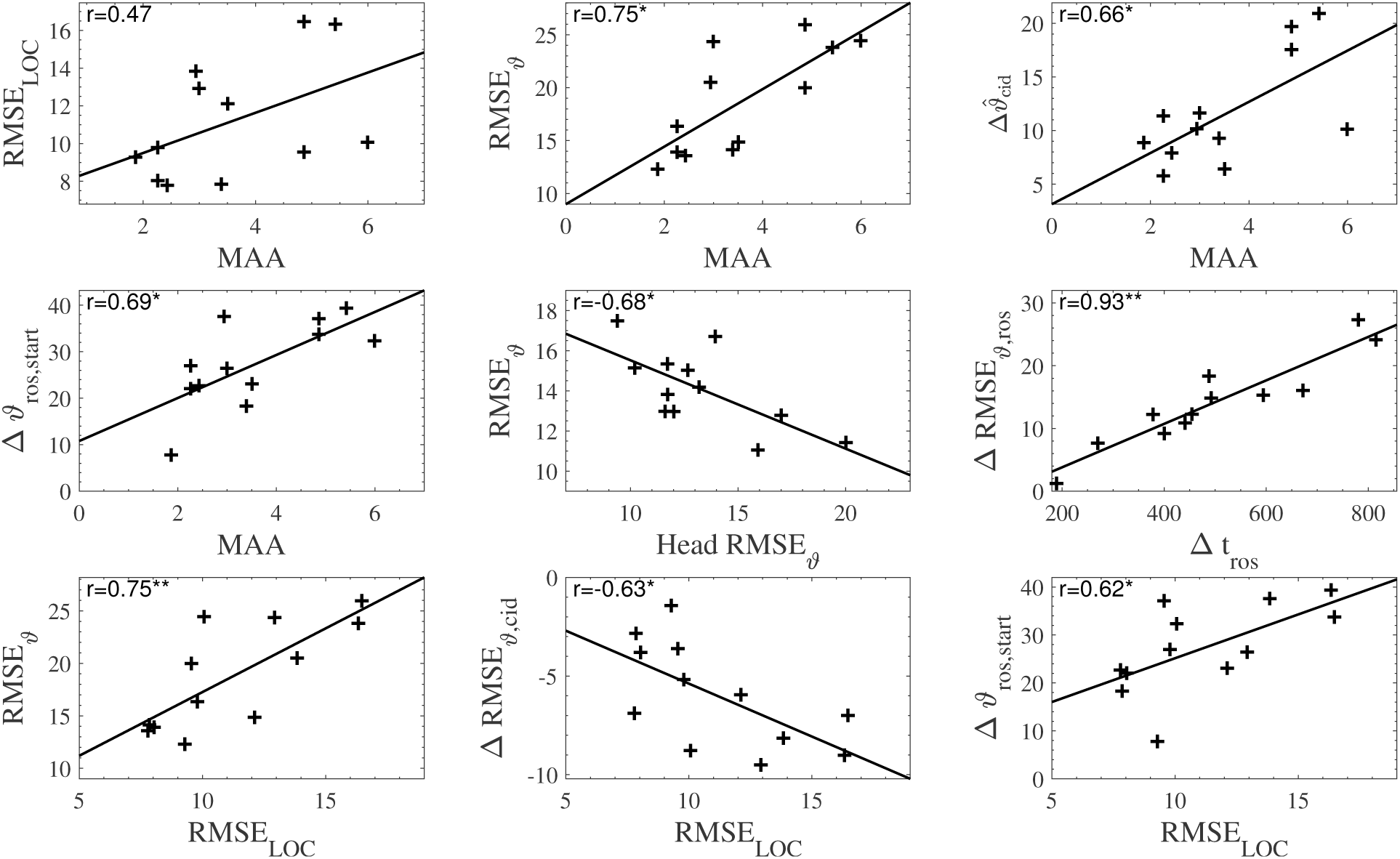
Scatter plots of selected outcome measures obtained from the sound localization, discrimination and tracking tests. For each plot, the Pearson correlation coefficient (r) is shown (statistically significant correlations are indicated by asterisks). Data from dynamic tests refer to the alternating trajectory with GUI, except for the data for the correlation involving head tracking, which refers to the steady trajectory. The outcome measures shown were averaged for each subject: minimum audible angle (MAA), root mean square error of the static localization test (RMSE_LOC_), position tracking error (RMSE_*ϑ*_), angular offset after a CID 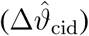, tracking error at the start of ROSs (Δ*ϑ*_*ros,start*_), head position tracking error (Head RMSE_*ϑ*_), position tracking error with GUI (RMSE_*ϑ*_),improvement after CID (ΔRMSE_*ϑ*,CID_), and duration of ROSs with GUI (Δ*t*_ros_). * p<.05, **p<.01

The absolute sound localization accuracy (indicated by RMSE_LOC_) showed a weak tendency to correlate with the MAA performance of each subject. In addition, we found significant correlations with the RMSE_LOC_ and sound tracking with many CIDs, as in the alternating trajectory. Subjects with better absolute sound localization accuracy performed better during sound tracking. Furthermore, the static localization ability correlated with the re-localization ability after a CID (see Figure 8) and the offset at the beginning of a source search 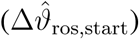. In addition, the duration of a source search Δt_ros_ (ms), was closely related to the improvement 6RMSE_*ϑ*,ros_ after the search.

Not only absolute but also relative localization performance (MAA) significantly correlated with sound tracking accuracy (RMSE_*ϑ*_) for a subject. Moreover, the MAA had a statistically significant correlation with two additional metrics assessed during sound tracking experiments: the absolute offset at a search start (Δ*ϑ*_ros,start_) and the angular offset after a CID 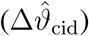.

We further found an indication that the more exactly subjects followed the source with their head in the dynamic case, the worse they performed in the dynamic test condition with restricted head movements.

In addition, we analyzed the correlation between sound localization and discrimination as in (Recanzone et al. 1998). Each data pair consisted of the standard deviation for a measurement location in the static sound localization test and the corresponding 68% MAA value. This analysis revealed a significant (p<0.01) but weak correlation (r=0.43) for the matching measurement positions between the tests (0°, 90°, 270° and 360°).

## 4 Discussion

Although relevant outcomes can be obtained, the assessment of spatial hearing performance remains underrepresented in clinical practice. The main reasons are the need for dedicated, expensive measurement setups and the time-consuming test procedures. In this context, most studies have evaluated sound localization performance in the frontal horizontal hemisphere with static sound scenarios (Grantham et al. 2007; Kerber & Seeber 2012; Jones et al. 2014; Dorman et al. 2016). In the future, modern hearing aid and implant audio processors will apply advanced, data science-based auditory scene analysis for improved hearing outcomes (Ma et al. 2017; Adavanne et al. 2019). Since users will rely on these technologies on a daily basis, reliable audiological validation is required. Our presented system is a first step in this direction because physically moving sound sources are robust against room characteristics or the head position of the listener (Zhang et al. 2017; Nelson et al. 2019; ISO 2009). Indeed, Ludwig et al. (2019) showed that children with auditory processing disorders performed normally under more realistic sound field localization tasks in contrast to headphone tests. The system can also be seen as an intermediate step towards virtual acoustics test systems. The presented approach could be useful to provide labelled reference data in dynamic settings with multiple independent sound sources under realistic conditions. The easy and reliable creation of ground truth data could be further beneficial for audio signal processor development or computational auditory scene analysis.

The major limitation of our system is the presence of movement noise of the speakers. In the tests with moving speakers, we mitigated this limitation by either providing masking movements (MAA test) or playing the stimulus from the single, moving speaker (tracking tests). Future versions of the WCARs could be driven by linear DC motors guided with a plain bearing. Such a drive system allows a very precise and rapid displacement of sound sources without motor noise and with minimal friction noise. The presented robot-based measurement setup provides reliable results of complex auditory measurements while being easy to use in common clinical environments. In the following section, our specific results are discussed.

### 4.1 Sound source localization

Our results are in line with those of the localization study of Makous & Middlebrooks (1990). They reported that the majority of errors were smaller than 5° in the frontal and smaller than 10° in the dorsal azimuths. We also observed lower errors in the frontal azimuth than in the dorsal azimuth; however, our errors were generally larger. One reason for this could have been the anechoic conditions in Makous & Middlebrooks (1990), which reduce localization errors (Giguère & Abel 1993).

We observed a larger RMSE_LOC_ on the left side than on the right side. As we did not measure any left-handed persons, the offset to the left side was consistent with the findings in Ocklenburg et al. (2010), where a hand-pointing method was used and participants showed a bias in sound localization to the side contralateral to the preferred hand. As we counted the GUI input as a hand-pointing method, the findings of Lewald et al. (2000) are relevant for error evaluation. These authors stated that hand pointing resulted in overshooting responses for targets in the frontal azimuths. However, most of our subjects jumped to the rough position on the GUI and afterwards made little adjustments. Response time with our GUI did not differ across directions and had no influence on the localization performance. A comparable setup with hidden loudspeakers, pink noise cues and a GUI as a feedback device was used in Jones et al. (2014). They reported an RMSE_LOC_ for monosyllabic words spoken by a male talker 8.2° in the frontal azimuths, including the sides. This finding confirms our results, as we measured an RMSE_LOC_ of 10.3°within this area. Studies not using a GUI resulted in RMSEs between 5.1°to 6.7°for a setup spanning −80°to 80° (Grantham et al. 2007; Kerber & Seeber 2012; Dorman et al. 2016). Only a few studies performed measurements within the full azimuthal plane. In addition to more comprehensive insight into the localization ability of the subjects, this approach enables the evaluation of FBCs. For broadband noise stimuli in the free field, Makous & Middlebrooks (1990) found average rates of FBCs for 6% of the stimuli, Pastore et al. (2018) found 5% and Best et al. (2010) reported FBC occurrences of 5%. In this study, we observed FBCs in only 3.6% of the stimuli.

Our setup overcame the bias introduced by prior knowledge of the sound locations Carlile et al. 1997. This effect was further reduced by the continuous feedback possibility. We preferred this feedback method to pointing via head or body rotations, as used in Makous & Middlebrooks (1990), due to the minor involvement of motor tasks. The trade off between mental effort when transferring the internal head-centered coordinate system on a GUI and the motor task effort with head or body rotations on measuring localization performance remains open. In Ocklenburg et al. (2010), no significant difference between hand and head pointing was observed for localization in the frontal azimuth. In contrast to the head pointing method in Makous & Middlebrooks (1990), our response time was twice as high and constant across directions.

### 4.2 Sound source discrimination

For the time-consuming MAA, we used a robot-supported, automated test procedure that acquired free-field MAAs in the full azimuthal plane at every 45°position (i.e., 8 tested directions in total) within 40 minutes, with 5 minutes to reliably assess the MAA at a given azimuth. Especially with multiple real sound sources at multiple azimuths, it is a difficult test to perform (Akeroyd & Whitmer 2016). Our results confirmed that the MAA increases with increasing displacement from the forward direction to the sides (Mills 1958). Furthermore, we confirmed the statement in Blauert (1997, p.41) that the localization blur in the back is approximately twice the value for the front. We further reproduced the results from Senn et al. (2005) and Mantokoudis et al. (2011) except for the increasing MAA at 180° in Senn et al. (2005). There is sparse literature on MAA results in the sound field at dorsal azimuths; therefore, our finding remains open to further discussion. However, as all our static and dynamic localization tests as well as those of Blauert (1997, p. 41) and Mantokoudis et al. (2011) showed good performance at 180°, we question the result of Senn et al. (2005) at this specific measurement location.

The literature contains two opinions regarding the relation between relative and absolute sound localization performance in humans. We were able to confirm the finding of Recanzone et al. (1998) that relative localization (test similar to MAA) thresholds at specific directions in the frontal plane are a poor indicator for the absolute localization performance in these directions. This finding is contradictory to that of Hartmann & Rakerd (1989), who stated that the MAA is an absolute rather than a relative localization task. In contrast to Freigang et al. (2014), we did not find a correlation in the frontal azimuth but did find a significant correlation in the dorsal azimuth^15^. Recanzone et al. (1998), claim that MAA thresholds are limited by absolute sound localization abilities. A significant correlation between the RMSE_LOC_ of the static localization test and the overall MAA was not observed in our study, but a trend was visible in the scatter plot (see Figure 10). An analysis between absolute and relative sound localization as in Recanzone et al. (1998), revealed a significant but weak correlation (r=0.43).

### 4.3 Sound source tracking

Our results demonstrate the validity of the tracking tests with real dynamic sound sources in the sound field. A MAA measurement at all positions took approximately 40 minutes in our study. The dynamic test of the alternating trajectory took approximately 5 minutes. Due to the correlation of both tests, we propose that a 5-minute single dynamic tracking test could potentially provide a comparable amount of information as the full MAA test battery, representing an 8-fold time savings. In addition to the correlation with sound discrimination abilities, we observed correlations of the tracking test with absolute localization performance. Because azimuthal sound motion detection and processing were directly connected to static source localization and discrimination abilities, our data support the snapshot theory of auditory motion perception for slow velocities, as described in Grantham (1997). The close relationship between the minimum audible movement angle (MAMA) and the MAA is obvious. The MAMA is 2-3 times larger than its corresponding MAA when measured under comparable conditions (Carlile & Leung 2016). The results of our dynamic source tracking tests with a GUI confirmed the results of this statistical relationship for the alternating trajectory. We observed a strong correlation with a slope of 2.7 between the RMSE_*ϑ*_ and the MAA values. In comparing the MAA results and the GUI-based results, similarities are evident. Both results show the smallest errors at 0° and 180° and have larger errors in the dorsal than in the frontal hemifield.

The tracking performance was worst when entering the dorsal azimuth or at a CID on the sides. This finding is in line with poor MAA results at the sides. It would be of great interest to investigate a direction-dependent MAA to differentiate between clockwise or counterclockwise stimuli shifts. As in the MAA test, the tracking in the frontal azimuth (±45°) was particularly good. An explanation could be that this area lies within the field of view. Training sound-tracking skills in everyday life is possible in this area. This could also explain the poor performance when entering the dorsal azimuth. We observed significant correlations between the static discrimination abilities of sound sources in terms of MAAs and the errors in the tracking tests, which forced a re-localization of the sound source. The MAA correlated strongly with the angular offset after a CID, 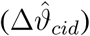. Furthermore, regardless of CIDs, an error at the beginning of a search correlated significantly with the average MAA performance. In the context of auditory motion perception, these findings support the snapshot theory (Grantham 1997).

In Moua et al. (2019), listeners were asked to distinguish between broadband noise (of 500, 1000 or 2000 ms durations) that appeared to be moving vs stationary in the frontal horizontal plane. The trajectories (0°to 40°) were travelled at different speeds depending on the duration of the stimulus. As this paper focuses on free-field tests, it must be mentioned that the test was based on non-individualized binaural recordings. Their results suggest that stationary sound source localization is better than the start- or end-point localization of a moving sound source. Within this study, we observed a strong correlation between the performance of static localization RMSE_LOC_ and the localization of a moving sound source RMSE_*ϑ*_ for the alternating trajectory. After the CIDs, we observed that the lower the RMSE_LOC_, the more accurately the sound was relocalized. In addition to CIDs, the size of the error at which a re-localization of a sound source was started by the subjects correlated significantly with the RMSE_LOC_. As with the MAA, this correlation supports the snapshot theory (Grantham 1997).

Subjects in Moua et al. (2019) were able to identify the direction of the sound source with an accuracy of >80%. The longer the angular distance of the trajectory was, the better the results. Our study with the steady trajectory confirmed this finding. When introducing CIDs as in the alternating trajectory, short source searches (∼ 500ms) appeared, but the accuracy of the correct movement was still higher than 80%. We observed that the longer the search time was, the higher the improvement of the localization error at the end of the search. Interestingly, the search time did not correlate with the size of the error at the beginning of the search. For a CID at the 45°position (appearing in both trajectories), the reaction time was 28% higher after a long ongoing movement, as in the steady trajectory, compared to the shorter movements of the alternating trajectory. Within subjects, the reaction times at this position correlated significantly between the steady and alternating trajectories (p<0.05). The scatter plot in the supplementary materials shows a strong linear trend^16^. This finding was in line with that of Moua et al. (2019), where the RMSE for end-point estimation of a moving sound source was lowest for the shortest trajectory. We hypothesize that this behavior can be explained by a model combining the initial belief (the prior) with the new observation (the likelihood) (Clark 2013). The prior, which refers to the belief that the movement continues as before, is higher for longer movement durations and thus has a greater influence on perception.

We observed a high tendency (>80%) of users to track the sound source ahead of its actual position. As with the reaction time for the steady trajectory, we hypothesize that subjects used previous experience to predict what they would hear. For smooth-pursuit eye movements, this effect was observed by Barnes (2008).

The point behind the head (180°) played a special role in the dynamic tracking of a sound source. As in our static localization and discrimination tests, a particularly good performance at 180°could be determined, although adjacent azimuths showed poor localization, discrimination and tracking performance. This finding again supports the snapshot theory of auditory motion perception (Grantham 1997). For both the steady and the alternating trajectories, RMSE_*ϑ*_ decreased to local minima when approaching the 180°position (see Figures 5 and 6). To the best of our knowledge, there was no study or setup that could measure this effect with real sound sources. Only at the back did the tracking accuracy increase after the CID (see Figure 8). Previous studies showed a dependency of tracking abilities on the azimuthal position and angular velocity of the moving sound source (Perrott & Musicant 1977; Chandler & Grantham 1992; Schmiedchen et al. 2013; Feinkohl et al. 2014). Therefore, the phenomena observed and described above should be further investigated with different stimulus velocities.

Current research tends to state that for auditory motion detection, current locations at different times are combined and weighted rather than the presence of motion sensitivity systems in the auditory system being considered (Grantham 1997; Carlile & Leung 2016). This statement includes the notion that absolute and relative sound localization abilities are indispensable in tracking and localizing moving sound sources. We observed correlations with sound localization and discrimination abilities and tracking performance, which supported this statement.

### 4.4 Gaze and dynamic tracking of sound sources

Head movement significantly improved the tracking performance of moving sounds in the free field. However, we observed higher tracking errors in gaze trajectories than in head trajectories. This finding is remarkable, as we did not use a head-pointing method (see above) but compared two conditions. In the first condition, head movements were restricted, and in the second condition, head movements were allowed. Furthermore, performance differed significantly between the GUI and the gaze-based input on our dynamic tracking tests. To the best of our knowledge, there exists no comparison of gaze versus GUI input for tracking real moving sound sources. The analysis of the head movement trajectories in this study showed a medium correlation. The more the additional acoustic information by head movements was used, the more difficult it was to track the sound without this information (see Figure 10). The correlation applied only for the steady trajectory, where continuous head movements were beneficial. The results were consistent with the idea discussed in Carlile & Leung (2016) that feedback from head motor commands or vestibular information may influence auditory perception. In addition, these results might show the tendency of a learned dependence on head movement to sound source tracking.

For static sound localization tests, using gaze-based input measures is an established and feasible methodology (Grantham et al. 2007; Volck et al. 2015). In Zambarbieri (2002), an error of 3°compared to visual target localization was found for measurements in the azimuth ± 35°. This error lies within auditory localization errors (Grantham et al. 2007; Kerber & Seeber 2012; Dorman et al. 2016). In contrast to saccades, smooth-pursuit eye movements cannot be controlled voluntarily. They require moving visual signals to be followed, auditory cues are not sufficient (Boucher et al. 2004; Berryhill et al. 2006). However, saccades can be evoked by auditory targets because they provide a position reference signal in space. Our results showed that head orientation was a more accurate estimator for the position of a moving stimulus than gaze positions (p<0.05). This result is remarkable in that head movement was not mandatory but optional when tracking the sound source. One reason for the better performance in contrast to gaze positions was certainly that continuous head movements without visual stimulation are possible in contrast to saccades. Inspection of the data revealed that the majority of errors came from the discrete angle due to the nature of saccades. We observed the majority of the saccades to be smaller than 10°, with the highest occurrences in the bin ranging from 0°to 4°. A histogram of the saccade sizes for the comparison with the GUI can be found in the supplementary materials^17^. Most saccade sizes correspond to the distance of the fiducial markers placed on the sound-transparent curtain in front of the subject (see Figure 1). In addition to the stepsize of the saccades, large errors can accidentally occur when gaze data are constantly recorded for dynamic tests. For a very accurate dataset, a higher commitment to the task than with the GUI is necessary. Tests to ensure oculomotor function could further reduce possible sources of inaccuracy (DiCesare et al. 2017). A small contribution to the error came from calibration accuracy, which was 2.6° ± 1.0°. In addition to the drawbacks mentioned above, eye-tracking allows the independent investigation of the impact of head movements on localization in dynamic test environments. We measured a significant improvement in tracking accuracy when head movement was allowed. To further improve the accuracy of gaze-based feedback in dynamic settings, one should decrease the distance of optical landmarks to reduce saccade stepsize.

## 5 Conclusion

The presented system and study enabled a comprehensive examination of the sound source localization, discrimination and tracking performance of normal-hearing subjects. The results of previous studies were confirmed and put in the context of novel findings from source tracking experiments. Our data supported the snapshot theory for slow stimulus velocities (Grantham 1997), as we observed correlations between the sound localization discrimination and tracking abilities. The largest tracking errors occurred when the stimulus crossed the interaural axis towards the dorsal azimuth. However, when approaching 180°, performance increased again. Furthermore, we observed a significant improvement in the tracking of a sound source if head movement was no longer restricted. However, gaze positions should be used with caution as feedback possibility for sound-tracking experiments in the free field. We observed that saccades introduced a higher tracking inaccuracy than a GUI on a touchpad device. Future studies will employ the test system to obtain the sound source localization, discrimination and tracking abilities of subjects with hearing impairment and cochlear implants

## Supporting information

SuppPub1

## Funding

This study was in part funded by the Med-El Corporation, the Berne University Research Foundation and the Fondation Charidu.

## Conflict of interest

The authors have no conflicts of interest to disclose.

## Acknowledgments

We would like to thank Dr. Thomas Wyss for his support with the integration of the robotic system.

## Supplemental Digital Content

- Supplemental Data (Figures, Tables, Screenshots)
  - SuppPub1.pdf
- Supplemental Data (Videos showing dynamic test procedures)
  - MM1.avi
  - MM2.avi
  - MM6.avi
  - MM4.avi
  - MM5.avi
  - MM6.avi

## Footnotes

1 See Table 1 in the supplementary materials at SuppPub1.pdf for a detailed overview of the demography.

2 See Figures 1 and 2 in the supplementary materials at SuppPub1.pdf for a more detailed description regarding the movement noise of the WCARs.

3 See Figure 6 in the supplementary materials at SuppPub1.pdf for an illustration of the GUI during an MAA test.

4 See Figure 8 in the supplementary materials at SuppPub1.pdf for an illustration of the GUI. The Figure shows the dial for the tracking test as well as for the static sound localization test.

5 See Tables 2-4 in the supplementary materials at SuppPub1.pdf for a detailed, tabular description of the trial and GUI test trajectories.

6 Video of the trial trajectory performed by a WCAR during a sound source tracking with touch pad test. MM1.avi (0.2MB)

7 Video of the steady trajectory performed by a WCAR during a sound source tracking with touch pad test. MM2.avi (1.6MB)

8 Video of the alternating trajectory performed by a WCAR during a sound source tracking with touch pad test. MM3.avi (3.6MB)

9 See Tables 5-7 in the supplementary materials at SuppPub1.pdf for a detailed, tabular description of the trial and test trajectories of the sound source tracking with/without head movement test.

10 Video of the trial trajectory performed by a WCAR during a sound source tracking with/without head movement test. MM4.avi (0.1MB)

11 Video of the steady trajectory performed by a WCAR during a sound source tracking with/without head movement test. MM5.avi (0.3MB)

12 Video of the alternating trajectory performed by a WCAR during a sound source tracking with/without head movement test. MM6.avi (0.3MB)

13 See Figures 3 and 4 in the supplementary materials at SuppPub1.pdf for overall RMSE_LOC_ results and direction specific RMSE_LOC_ values for each subject.

14 See Figures 5 and 7 in the supplementary materials at SuppPub1.pdf for overall MAA values and direction specific MAA values for each subject.

15 See Figure 10 in the supplementary materials at SuppPub1.pdf for the scatter plot between the MAA at the dorsal positions and the static localization error (RMSE_LOC_) at the dorsal positions.

16 See Figure 11 in the supplementary materials at SuppPub1.pdf for the scatter plot between the reaction times Δt_cid_ in both the steady and alternating trajectory. Measurement position was the CID at 45° measurement position.

17 See Figure 9 in the supplementary materials at SuppPub1.pdf for a histogram of the saccade sizes in the GUI vs Gaze-based input comparison.

## References

Adavanne, S., A. Politis, T. Virtanen (2019). Localization, Detection and Tracking of Multiple Moving Sound Sources with a Convolutional Recurrent Neural Network. arXiv: 1904.12769.

Akeroyd, M. A., W. M. Whitmer (2016). Spatial Hearing and Hearing Aids. In: 181–215. DOI: 10.1007/978-3-319-33036-5_7.

Barnes, G. (2008). Cognitive processes involved in smooth pursuit eye movements. Brain Cogn, 68, 309–326. DOI: 10.1016/J.BANDC.2008.08.020.

Berens, P. (2009). CircStat: A MATLAB Toolbox for Circular Statistics. J Stat Softw, 31, 1–21. DOI: 10.18637/jss.v031.i10.

Berryhill, M. E., T. Chiu, H. C. Hughes (2006). Smooth Pursuit of Nonvisual Motion. J Neurophysiol, 96, 461–465. DOI: 10.1152/jn.00152.2006.

Best, V., S. Kalluri, S. McLachlan, et al. (2010). A comparison of CIC and BTE hearing aids for three-dimensional localization of speech. Int J Audiol, 49, 723–732. DOI: 10.3109/14992027.2010.484827.

Blauert, J. (1997). Spatial hearing: the psychophysics of human sound localization. Cambridge, MA: MIT Press.

Boucher, L., A. Lee, Y. E. Cohen, et al. (2004). Ocular tracking as a measure of auditory motion perception. J Physiol Paris, 98, 235–248. DOI: 10.1016/j.jphysparis.2004.03.010.

Brimijoin, W. O., M. A. Akeroyd (2014). The moving minimum audible angle is smaller during self motion than during source motion. Front Neurosci, 8, 273. DOI: 10.3389/fnins.2014.00273.

Carlile, S., P. Leong, S. Hyams (1997). The nature and distribution of errors in sound localization by human listeners. Hear Res, 114, 179–196. DOI: 10.1016/S0378-5955(97)00161-5.

Carlile, S., J. Leung (2016). The Perception of Auditory Motion. Trends Hear, 20. DOI: 10.1177/2331216516644254.

Chandler, D. W., D. W. Grantham (1992). Minimum audible movement angle in the horizontal plane as a function of stimulus frequency and bandwidth, source azimuth, and velocity. J Acoust Soc Am, 91, 1624–1636. DOI: 10.1121/1.402443.

Cherry, E. C. (1953). Some Experiments on the Recognition of Speech, with One and with Two Ears. The Journal of the Acoustical Society of America, 25, 975–979. DOI: 10.1121/1.1907229.

Clark, A. (2013). Whatever next? Predictive brains, situated agents, and the future of cognitive science. Behavioral and Brain Sciences, 36, 181–204. DOI: 10.1017/S0140525X12000477.

Denk, F., S. D. Ewert, B. Kollmeier (2019). On the limitations of sound localization with hearing devices. The Journal of the Acoustical Society of America, 146, 1732–1744. DOI: 10.1121/1.5126521.

DiCesare, C. A., A. W. Kiefer, P. Nalepka, et al. (2017). Quantification and analysis of saccadic and smooth pursuit eye movements and fixations to detect oculomotor deficits. Behav Res Methods, 49, 258–266.

Dorman, M. F., L. H. Loiselle, S. J. Cook, et al. (2016). Sound source localization by normal hearing listeners, hearing-impaired listeners and cochlear implant listeners. Audiol Neurootol, 21, 127. DOI: 10.1159/000444740.

Feinkohl, A., S. M. Locke, J. Leung, et al. (2014). The effect of velocity on auditory representational momentum. J Acoust Soc Am, 136, EL20–EL25. DOI: 10.1121/1.4881318.

Freigang, C., K. Schmiedchen, I. Nitsche, et al. (2014). Free-field study on auditory localization and discrimination performance in older adults. Exp Brain Res, 232, 1157–1172. DOI: 10.1007/s00221-014-3825-0.

Geronazzo, M., E. Peruch, F. Prandoni, et al. (2019). Applying a single-notch metric to image-guided head-related transfer function selection for improved vertical localization. J Audio Eng Soc, 67, 414–428.

Giguère, C., S. M. Abel (1993). Sound localization: Effects of reverberation time, speaker array, stimulus frequency, and stimulus rise/decay. The Journal of the Acoustical Society of America, 94, 769–776. DOI: 10.1121/1.408206.

Grantham, D. W. (1997). Auditory motion perception: Snapshots revisited. Binaural and spatial hearing in real and virtual environments, 295–313.

Grantham, D. W., D. H. Ashmead, T. A. Ricketts, et al. (2007). Horizontal-Plane Localization of Noise and Speech Signals by Postlingually Deafened Adults Fitted With Bilateral Cochlear Implants*. Ear Hear, 28, 524–541. DOI: 10.1097/AUD.0b013e31806dc21a.

Hartmann, W. M. (1983). Localization of sound in rooms. The Journal of the Acoustical Society of America, 74, 1380–1391. DOI: 10.1121/1.390163.

Hartmann, W. M., B. Rakerd (1989). On the minimum audible angle—A decision theory approach. J Acoust Soc Am, 85, 2031–2041. DOI: 10.1121/1.397855.

International Organization for Standardization (2009). Part 2: Sound field audiometry with pure-tone and narrow-band test signals. In: Acoustics – Audiometric test methods. ISO 8253-2:2009, 1–16.

Jones, H., A. Kan, R. Y. Litovsky (2014). Comparing Sound Localization Deficits in Bilateral Cochlear-Implant Users and Vocoder Simulations With Normal-Hearing Listeners. Trends Hear, 18. DOI: 10.1177/2331216514554574.

Kerber, S., B. U. Seeber (2012). Sound Localization in Noise by Normal-Hearing Listeners and Cochlear Implant Users. Ear Hear, 33, 445–457. DOI: 10.1097/AUD.0b013e318257607b.

Kourosh, S., D. R. Perrott (1990). Minimum audible movement angles as a function of sound source trajectory. J Acoust Soc Am, 83, 2639–2644.

Letowski, T., S. Letowski (2011). Localization Error: Accuracy and Precision of Auditory Localization. In: Advances in Sound Localization. Ed. by P. Strumillo. Rijeka, Croatia: IntechOpen. Chap. 4. DOI: 10.5772/15652.

Lewald, J., G. J. Dörrscheidt, W. H. Ehrenstein (2000). Sound localization with eccentric head position. Behav Brain Res, 108, 105–25.

Ludwig, A. A., M. Zeug, M. Schönwiesner, et al. (2019). Auditory localization accuracy and auditory spatial discrimination in children with auditory processing disorders. Hear Res, 377, 282–291.

Ma, N., T. May, G. J. Brown (2017). Exploiting Deep Neural Networks and Head Movements for Robust Binaural Localization of Multiple Sources in Reverberant Environments. IEEE/ACM Transactions on Audio, Speech, and Language Processing, 25, 2444–2453. DOI: 10.1109/TASLP.2017.2750760.

Makous, J. C., J. C. Middlebrooks (1990). Two-dimensional sound localization by human listeners. J Acoust Soc Am, 87, 2188–2200. DOI: 10.1121/1.399186.

Mantokoudis, G., M. Kompis, M. Vischer, et al. (2011). In-the-canal versus behind-the-ear microphones improve spatial discrimination on the side of the head in bilateral cochlear implant users. Otology and Neurotology, 32, 1–6. DOI: 10.1097/MAO.0b013e3182001cac.

Martinez, A. M. C., L. Gerlach, G. Payá-Vayá, et al. (2019). DNN-based performance measures for predicting error rates in automatic speech recognition and optimizing hearing aid parameters. Speech Commun, 106, 44–56.

Middlebrooks, J. C. (2015). Sound localization. Handb Clin Neurol, 129, 99–116. DOI: 10.1016/B978-0-444-62630-1.00006-8.

Mills, A. W. (1958). On the Minimum Audible Angle. J Acoust Soc Am, 30, 237–246. DOI: 10.1121/1.1909553.

Moua, K., A. Kan, H. G. Jones, et al. (2019). Auditory motion tracking ability of adults with normal hearing and with bilateral cochlear implants. J Acoust Soc Am, 145, 2498–2511. DOI: 10.1121/1.5094775.

Mueller, M. F., A. Kegel, S. M. Schimmel, et al. (2012). Localization of virtual sound sources with bilateral hearing aids in realistic acoustical scenes. J Acoust Soc Am, 131, 4732–4742.

Naithani, G., T. Barker, G. Parascandolo, et al. (2017). Low latency sound source separation using convolutional recurrent neural networks. In: 2017 IEEE Workshop on Applications of Signal Processing to Audio and Acoustics (WASPAA). IEEE. New Paltz, NY, 71–75.

Nelson, E., R. M. Reeder, L. K. Holden, et al. (2019). Front-and rear-facing horizontal sound localization results in adults with unilateral hearing loss and normal hearing. Hear Res, 372, 3–9.

Ocklenburg, S., M. Hirnstein, M. Hausmann, et al. (2010). Auditory space perception in left- and right-handers. Brain Cogn, 72, 210–217. DOI: 10.1016/j.bandc.2009.08.013.

Pastore, M. T., S. J. Natale, W. A. Yost, et al. (2018). Head Movements Allow Listeners Bilaterally Implanted With Cochlear Implants to Resolve Front-Back Confusions. Ear Hear, 39, 1224–1231. DOI: 10.1097/AUD.0000000000000581.

Perrott, D. R., A. D. Musicant (1977). Minimum auditory movement angle: Binaural localization of moving sound sources. J Acoust Soc Am, 62, 1463. DOI: 10.1121/1.381675.

Prepelită, S., M. Geronazzo, F. Avanzini, et al. (2016). Influence of voxelization on finite difference time domain simulations of head-related transfer functions. J Acoust Soc Am, 139, 2489–2504.

Razavi, B., W. E. O’Neill, G. D. Paige (2007). Auditory Spatial Perception Dynamically Realigns with Changing Eye Position. J Neurosci, 27, 10249–10258. DOI: 10.1523/JNEUROSCI.0938-07.2007.

Recanzone, G. H., S. D. D. R. Makhamra, D. C. Guard (1998). Comparison of relative and absolute sound localization ability in humans. J Acoust Soc Am, 103, 1085–1097. DOI: 10.1121/1.421222.

Romero-Ramirez, F. J., R. Muñoz-Salinas, R. Medina-Carnicer (2018). Speeded up detection of squared fiducial markers. Image Vis Comput, 76, 38–47. DOI: 10.1016/j.imavis.2018.05.004.

Schmiedchen, K., C. Freigang, R. Rübsamen, et al. (2013). A comparison of visual and auditory representational momentum in spatial tasks. Atten Percept Psychophys, 75, 1507–1519. DOI: 10.3758/s13414-013-0495-0.

Senn, P., M. Kompis, M. Vischer, et al. (2005). Minimum Audible Angle, Just Noticeable Interaural Differences and Speech Intelligibility with Bilateral Cochlear Implants Using Clinical Speech Processors. Audiol Neurootol, 10, 342–352. DOI: 10.1159/000087351.

Shelton, J., G. P. Kumar, J. Shelton, et al. (2010). Comparison between Auditory and Visual Simple Reaction Times. Neurosci Med, 01. DOI: 10.4236/NM.2010.11004.

Shen, Y., W. Dai, V. M. Richards (2015). A MATLAB toolbox for the efficient estimation of the psychometric function using the updated maximum-likelihood adaptive procedure. Behav Res Methods, 47, 13–26. DOI: 10.3758/s13428-014-0450-6.

Trattler, B., P. K. Kaiser, N. J. Friedman (2012). Review of ophthalmology. Philadelphia, PA: Saunders Elsevier, 255.

Van den Bogaert, T., T. J. Klasen, M. Moonen, et al. (2006). Horizontal localization with bilateral hearing aids: Without is better than with. J Acoust Soc Am, 119, 515–526.

Virtanen, T., M. D. Plumbley, D. Ellis, eds. (2018). Computational Analysis of Sound Scenes and Events. Cham: Springer International Publishing. DOI: 10.1007/978-3-319-63450-0.

Volck, A. C., R. D. Laske, R. Litschel, et al. (2015). Sound localization measured by eye-tracking. Int J Audiol, 54, 976–983. DOI: 10.3109/14992027.2015.1088968.

Wimmer, W., M. Kompis, C. Stieger, et al. (2017). Directional microphone contralateral routing of signals in cochlear implant users: A within-subjects comparison. Ear Hear, 38, 368–373. DOI: 10.1097/AUD.0000000000000412.

Winn, M. B., D. Wendt, T. Koelewijn, et al. (2018). Best Practices and Advice for Using Pupillometry to Measure Listening Effort: An Introduction for Those Who Want to Get Started. Trends Hear, 22, 1–32. DOI: 10.1177/2331216518800869.

Zambarbieri, D. (2002). The latency of saccades toward auditory targets in humans. Prog Brain Res, 140, 51–59. DOI: 10.1016/S0079-6123(02)40041-6.

Zhang, W., P. Samarasinghe, H. Chen, et al. (2017). Surround by sound: A review of spatial audio recording and reproduction. Appl Sci, 7, 532.

Zirn, S., J. Angermeier, S. Arndt, et al. (2019). Reducing the Device Delay Mismatch Can Improve Sound Localization in Bimodal Cochlear Implant or Hearing-Aid Users. Trends Hear, 23. DOI: 10.1177/2331216519843876.

