## Supplementary material for "Dynamic sound field audiometry: static and dynamic spatial hearing tests in the full horizontal plane": SuppPub1

### - SUPPLEMENTARY DIGITAL CONTENT -
### DYNAMIC SOUND FIELD AUDIOMETRY: SOUND SOURCE LOCALIZATION, DISCRIMINATION AND TRACKING TESTS WITH NORMAL HEARING SUBJECTS

Fischer T.[1,2], Kompis M.[1], Mantokoudis G.[1], Caversaccio M.[1,2], and Wimmer W.[1,2]

[1]Department of ENT, Head and Neck Surgery, Inselspital, Bern University Hospital, University of Bern, Bern 3008, Switzerland
[2]Hearing Research Laboratory, ARTORG Center for Biomedical Engineering Research, University of Bern, Bern 3008, Switzerland

October 21, 2019

#### 1 Demography

Table 1: Demography of the study population. All subjects were right-handed. PTA = Pure Tone Average, NH = Normal Hearing.

| Subject | PTA 4 ( left) | PTA 4 (right) | Sex | Age |
|---|---|---|---|---|
| NH01 | -6,3 | -6,3 | f | 31 |
| NH02 | 1,3 | -1,3 | m | 54 |
| NH03 | -3,8 | -8,8 | m | 25 |
| NH04 | 1,3 | 0,0 | m | 40 |
| NH05 | -5,0 | -5,0 | m | 42 |
| NH06 | -2,5 | -3,8 | f | 25 |
| NH07 | 1,3 | 6,3 | f | 44 |
| NH08 | -3,8 | -3,8 | m | 24 |
| NH09 | -7,5 | 0,0 | m | 28 |
| NH10 | -1,3 | 3,8 | m | 30 |
| NH11 | -2,5 | -2,5 | f | 27 |
| NH12 | -1,3 | -1,3 | m | 36 |

#### 2 Measurement setup

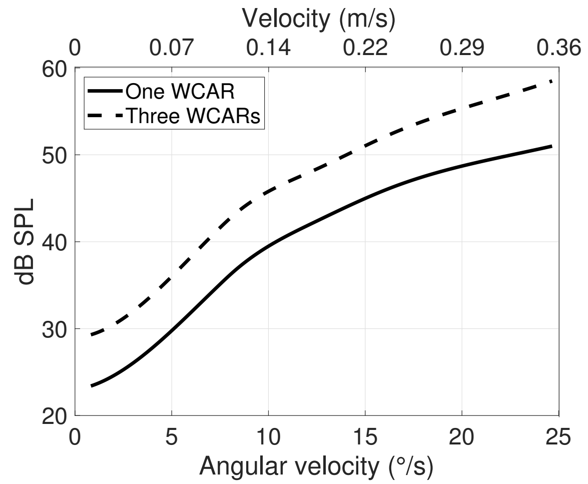


Figure 1: Movement noise of the Wireless Controllable Audio Robots (WCARs) at different angular velocities. The angular velocity used during the tracking tests was 7.4°/s.

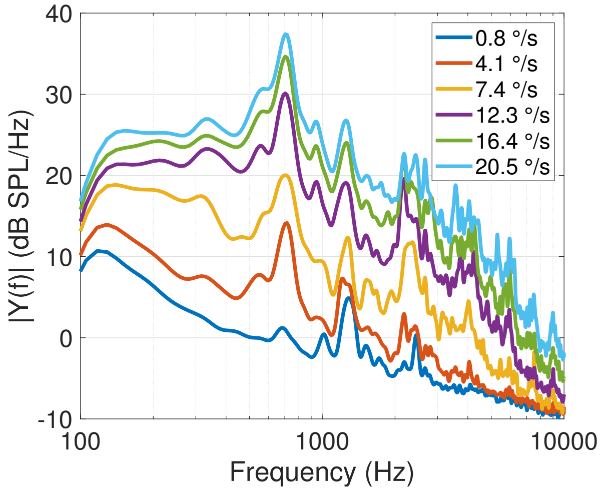


Figure 2: Fast Fourier transform of one moving Wireless Controllable Audio Robot (WCAR) at different velocities (1° corresponds to an arc length of 0.019 m). The angular velocity used during the tracking tests was 7.4°/s.

#### 3 Static sound source localization

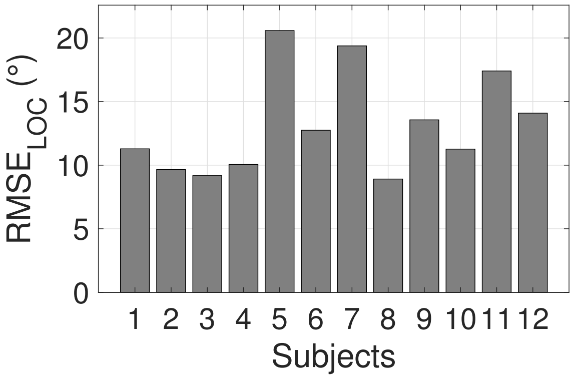


Figure 3: Static localization. Root mean square error values ($\mathrm{RMSE}_{\mathrm{LOC}}$) of each subject. Front-back confusions (FBCs) were excluded from the calculations.

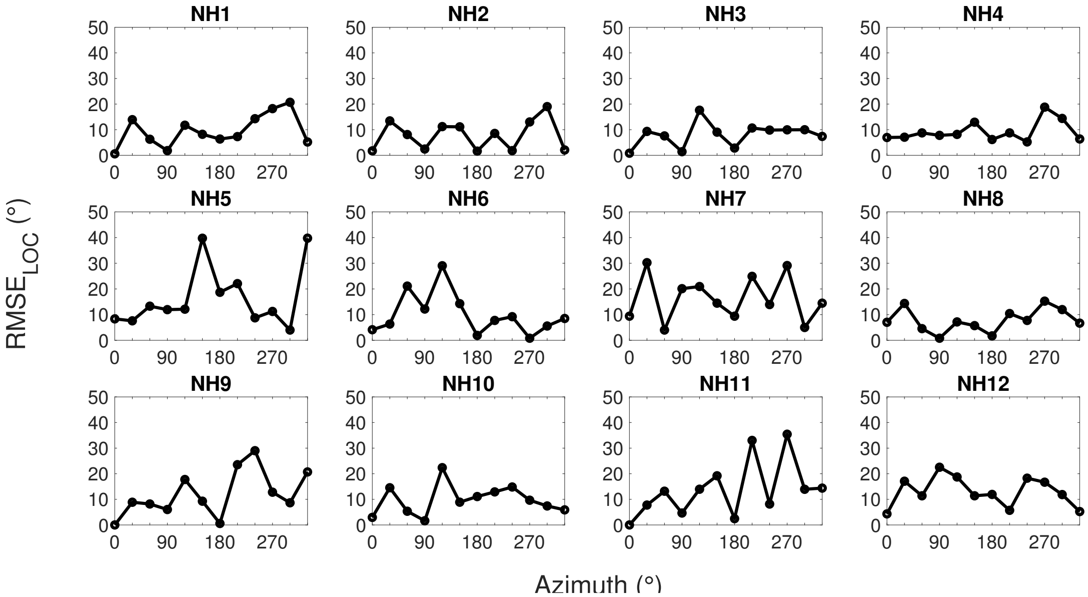


Figure 4: Static localization. Azimuth specific performance in linear plots for earch subject. Front-back confusions (FBCs) were excluded from the calculations.

#### 4 Minimum audible angle

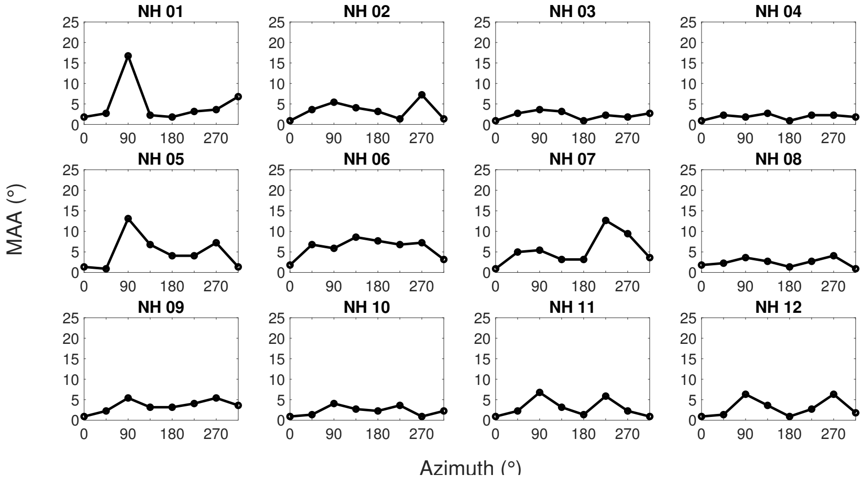


Figure 5: Minimum Audible Angle (MAA). Azimuth specific performance in linear plots for each subject.

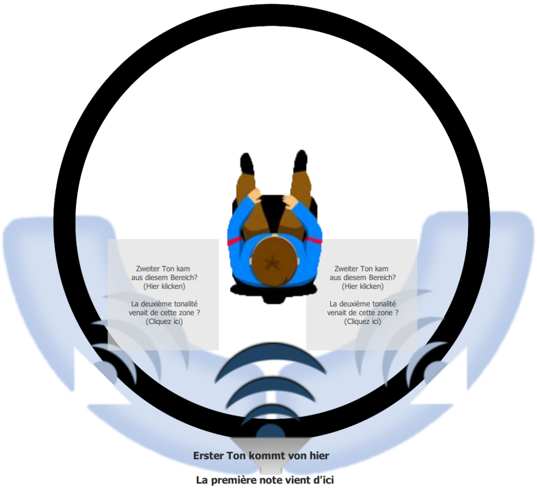


Figure 6: Touch-GUI for the Minimum Audible Angle (MAA) test (exemplary at 180°).

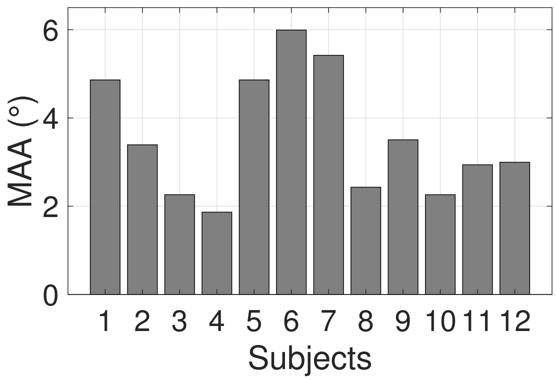


Figure 7: Azimuth unspecific mean value of the Minimum Audible Angle (MAA) for each subject.

#### 5 Sound source tracking with touch pad

Table 2: Trial trajectory of the GUI-based stimulus tracking.

| Start azimuth (°) | Movement direction | Travelled distance (°) | End azimuth (°) |
|---|---|---|---|
| 0 | Clockwise | 360 | 0 |
| 0 | Cntr.-Clockwise | 90 | 270 |

Table 3: Steady trajectory of the GUI-based stimulus tracking.

| Start azimuth (°) | Movement direction | Travelled distance (°) | End azimuth (°) |
|---|---|---|---|
| 315 | Clockwise | 450 | 45 |
| 45 | Cntr.-Clockwise | 450 | 315 |

Table 4: Alternating trajectory of the GUI-based stimulus tracking.

| **Start azimuth (°)** | **Movement direction** | **Travelled distance (°)** | **End azimuth (°)** |
|---|---|---|---|
| 0 | Clockwise | 45 | 45 |
| 45 | Cntr.-Clockwise | 45 | 0 |
| 0 | Clockwise | 90 | 90 |
| 90 | Cntr.-Clockwise | 45 | 45 |
| 45 | Clockwise | 90 | 135 |
| 135 | Cntr.-Clockwise | 45 | 90 |
| 90 | Clockwise | 90 | 180 |
| 180 | Cntr.-Clockwise | 45 | 135 |
| 135 | Clockwise | 90 | 225 |
| 225 | Cntr.-Clockwise | 45 | 180 |
| 180 | Clockwise | 90 | 270 |
| 270 | Cntr.-Clockwise | 45 | 225 |
| 225 | Clockwise | 90 | 315 |
| 315 | Cntr.-Clockwise | 45 | 270 |
| 270 | Clockwise | 90 | 0 |
| 0 | Cntr.-Clockwise | 45 | 315 |
| 315 | Clockwise | 45 | 0 |
| 0 | Cntr.-Clockwise | 90 | 270 |
| 270 | Clockwise | 45 | 315 |
| 315 | Cntr.-Clockwise | 90 | 225 |
| 225 | Clockwise | 45 | 270 |
| 270 | Cntr.-Clockwise | 90 | 180 |
| 180 | Clockwise | 45 | 225 |
| 225 | Cntr.-Clockwise | 90 | 135 |
| 135 | Clockwise | 45 | 180 |
| 180 | Cntr.-Clockwise | 90 | 90 |
| 90 | Clockwise | 45 | 135 |
| 135 | Cntr.-Clockwise | 90 | 45 |
| 45 | Clockwise | 45 | 90 |
| 90 | Cntr.-Clockwise | 90 | 0 |
| 0 | Clockwise | 45 | 45 |
| 45 | Cntr.-Clockwise | 45 | 0 |

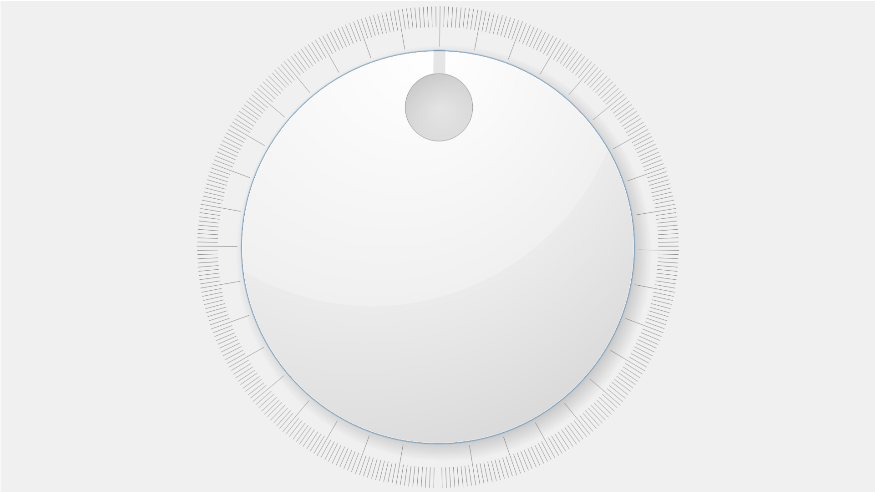

Figure 8: Touch-GUI for the tracking and static localization tests.

#### 6 Sound source tracking with eye movements or optional head movements

Table 5: Trial trajectory of the gaze- or head-based stimulus tracking.

| Start azimuth (°) | Movement direction | Travelled distance (°) | End azimuth (°) |
|---|---|---|---|
| 0 | Cntr.-Clockwise | 60 | 300 |
| 300 | Clockwise | 120 | 60 |
| 60 | Cntr.-Clockwise | 60 | 0 |

Table 6: Steady trajectory of the gaze- or head-based stimulus tracking.

| Start azimuth (°) | Movement direction | Travelled distance (°) | End azimuth (°) |
|---|---|---|---|
| 300 | Clockwise | 120 | 60 |
| 60 | Cntr.-Clockwise | 120 | 300 |
| 300 | Clockwise | 120 | 60 |
| 60 | Cntr.-Clockwise | 120 | 300 |
| 300 | Clockwise | 120 | 60 |
| 60 | Cntr.-Clockwise | 120 | 300 |

Table 7: Alternating trajectory of the gaze- or head-based stimulus tracking.

| Start azimuth (°) | Movement direction (°) | Travelled distance (°) | End azimuth (°) |
|---:|---|---:|---:|
| 300 | Clockwise | 15 | 315 |
| 315 | Cntr.-Clockwise | 15 | 300 |
| 300 | Clockwise | 15 | 315 |
| 315 | Cntr.-Clockwise | 15 | 300 |
| 300 | Clockwise | 60 | 0 |
| 0 | Cntr.-Clockwise | 45 | 315 |
| 315 | Clockwise | 45 | 0 |
| 0 | Cntr.-Clockwise | 45 | 315 |
| 315 | Clockwise | 90 | 45 |
| 45 | Cntr.-Clockwise | 45 | 0 |
| 0 | Clockwise | 45 | 45 |
| 45 | Cntr.-Clockwise | 45 | 0 |
| 0 | Clockwise | 60 | 60 |
| 60 | Cntr.-Clockwise | 15 | 45 |
| 45 | Clockwise | 15 | 60 |
| 60 | Cntr.-Clockwise | 15 | 45 |
| 45 | Clockwise | 15 | 60 |
| 60 | Cntr.-Clockwise | 105 | 315 |
| 315 | Clockwise | 90 | 45 |
| 45 | Cntr.-Clockwise | 90 | 315 |

##### 6.1 Comparison touch pad vs. gaze detection input

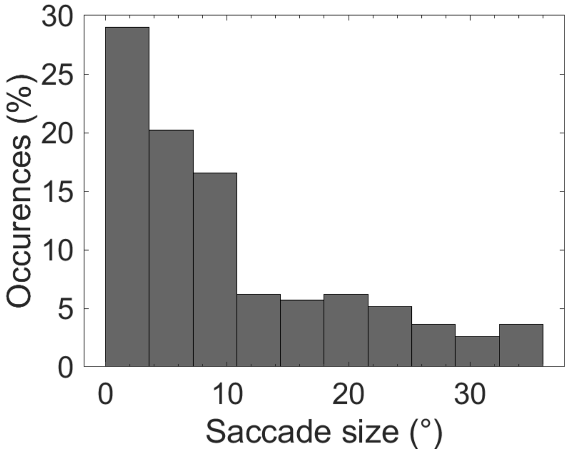


Figure 9: Saccade size for the dynamic tracking without head movement test (GUI vs Gaze-based input).

#### 7 Correlation analysis

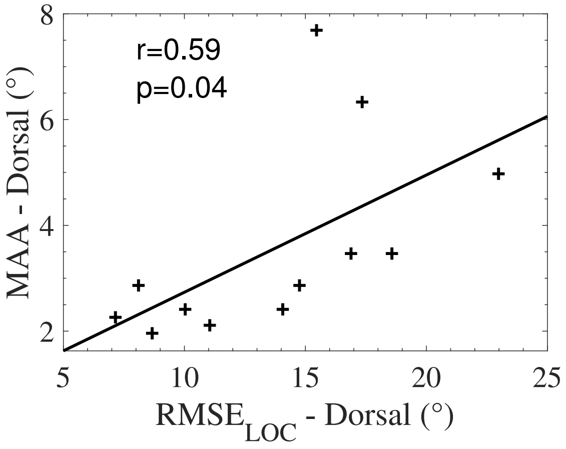


Figure 10: Minimum Audible Angle (MAA) at the dorsal positions (135°, 180°, 225°) and the static localization error ($\mathrm{RMSE_{LOC}}$) at the dorsal positions (120°, 150°, 180°, 210°, 240°).

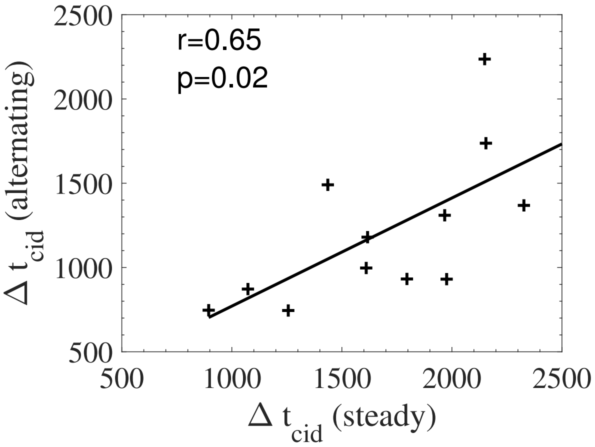


Figure 11: Scatter plot showing the reaction time $\Delta t_{cid}$ for a change in direction (CID) of the stimulus at 45° measurement position. The data refers to the GUI input method in the alternating or steady trajectory.
